# Planning and movement activities co-adapt in a motor-reference frame during 3D BCI-controlled reach adaptation in monkey frontal and parietal cortices

**DOI:** 10.1101/2024.11.03.621745

**Authors:** E. Ferrea, P. Morel, A. Gail

**Affiliations:** German Primate Center, Göttingen, Germany; Faculty of Biology and Psychology, University of Göttingen, Göttingen, Germany; Bernstein Center for Computational Neuroscience Göttingen, Göttingen, Germany; Leibniz ScienceCampus Primate Cognition, Göttingen, Germany; Institute for Neuromodulation and Neurotechnology, University Hospital and University of Tübingen, Tübingen, Germany; Univ. Littoral Côte d’Opale, Univ. Artois, Univ. Lille, ULR 7369-URePSSS-Unité de Recherche Pluridisciplinaire Sport Santé Société, F-62100 Calais, France

## Abstract

Perturbing visual feedback is a powerful tool for studying visuomotor adaptation. However, unperturbed proprioceptive signals in common paradigms inherently co-varies with physical movements and causes incongruency with the visual input. This can create challenges when interpreting underlying neurophysiological mechanisms. We employed a brain-computer interface (BCI) in rhesus monkeys to investigate spatial encoding in frontal and parietal areas during a 3D visuomotor rotation task where only visual feedback was movement-contingent. We found that both brain regions better reflected the adapted motor commands than the perturbed visual feedback during movement preparation and execution. This adaptive response was observed in both local and remote neurons, even when they did not directly contribute to the BCI input signals. The transfer of adaptive changes in planning activity to corresponding movement corrections was stronger in the frontal than in the parietal cortex. Our results suggest an integrated large-scale visuomotor adaptation mechanism in a motor-reference frame spanning across frontoparietal cortices.

## Introduction

Intracortical brain-computer interfaces (BCIs) have shown success in restoring lost motor functions in individuals with tetraplegia and chronic stroke, primarily by extracting signals from the motor cortex (1–6). Recent advancements have extended this approach to signals obtained from the parietal cortex (7,8). Besides their translational applications, BCI paradigms provide a crucial tool for studying the neurophysiological basis of motor learning (9–20).

Unlike conventional motor learning paradigms involving physical limb movement, BCI control establishes an experimentally controllable mapping from brain activity to effector motion through the neural decoder. Additionally, BCI paradigms help isolate the impact of visual feedback on movement adaptation by keeping proprioceptive feedback constant during visuomotor perturbations (16). This approach reduces potential confounds inherent in the adaptation of physical movements, which co-vary with proprioceptive feedback, and emphasizes the visual aspect of feedback control.

In our study, we employ a BCI visuomotor rotation paradigm in rhesus monkeys to explore the role of visual feedback and the contributions of parietal and premotor sensorimotor areas to BCI adaptation. We ask whether both brain regions (i) contribute to adaptation irrespective of an immediate impact on the produced motor behavior, (ii) reflect the same or different spatial dimensions of fast visuomotor adaptation, and (iii) show adaptation of motor planning activity consistent with motor behavior.

An open question about BCI learning is under which circumstances behavioral performance is improved by exploiting an existing repertoire of neural dynamic states (“within-manifold” learning) or by reconfiguring the neural network structure (“outside-manifold” learning). In response to perturbations of a BCI decoder, short-term learning does not appear to significantly affect the correlation patterns among neurons in the motor cortex (14,17,21). This suggests that at least the local network structure supplying input to the decoder (i.e., the controlling motor area) remains largely unchanged and constrains neural adaptation during short-term learning. However, with longer-term learning, the brain can be trained to generate new activity patterns in the controlling units that deviate from the expected correlation pattern of the network. Such outside-manifold learning aligns with the view that the tuning of individual neurons can be adapted independently during BCI learning (11,21–24).

A way how the decoder input could be adapted without changing the network structure is by means of “re-aiming” or “re-association”. According to this view, a specific neural activity pattern associated with compensating for the introduced visuomotor perturbation is recruited from the range of patterns available within the existing network structure (13,14,17,20). This view implies an unchanged network structure also for neighboring neurons, such that also non-controlling units in the controlling area that do not provide input to the decoder adapt their activity in a manner consistent with changes in the BCI controlling units. We hypothesize that the comparatively fast adaptation to visuomotor rotation, achievable within a single session of a few hundred trials, is accompanied by such re-association learning. We will test three implications of this hypothesis.

First, we will address the question whether changes in neural activity during BCI learning occur consistently not only in nearby non-controlling neurons, as shown previously (11,13,25), but also across a larger-scale sensorimotor network, including frontoparietal circuits. Previous studies in humans and monkeys have not extensively examined the extent of adaptation within a distributed neural network, including regions that are not directly involved in generating or updating motor commands but may still play a crucial role in sensorimotor integration. These regions, such as the parietal cortex, are expected to contribute to motor learning (26,27). In a human BCI motor learning study involving the prefrontal cortex, dorsal premotor cortex, primary motor and sensory cortices, and the posterior parietal cortex, neural changes occurred with long-term learning and were associated nonspecifically with reduced cognitive demand (28). In contrast, we aim to understand the mechanisms of fast adaptation, which is particularly desirable for BCI learning and, in conventional visuomotor rotation learning, is often associated with the updating of an internal model (29,30). We compare the contributions of non-controlling neurons residing locally next to the BCI controlling neurons in the frontal areas or remotely in parietal areas to test the hypotheses of re-association versus re-configuration. If the re-association hypothesis is true, we expect fast adaptation effects to generalize to remote brain regions.

Second, we ask whether premotor and parietal brain regions not only both adapt but do so in the same spatial frame of reference. Specifically, we test if neural changes reflect the adapted motor command or if the parietal cortex rather reflects the associated adapted sensory feedback. Posterior parietal networks integrate sensory information from multiple modalities, including visual input, and contain motor-goal information during reach planning without directly driving motor output (31,32). Preparatory activities include reach-goal information in different body-related frames of reference, including the predominantly gaze centered parietal reach region PRR. Preparatory activities include reach-goal information in different body-related frames of reference (33), which can be related to the visual and the physical goal of the movement (34), and the predominantly hand centered area 5d (35). Instead, dorsal premotor cortex encodes reach target locations in a more strongly mixed encoding of the relative positions of the hand, eye, and target (36,37), premotor and parietal areas encode the target location differently, with premotor cortex showing a more strongly mixed encoding of the relative positions of the hand, eye, and target (36,37)., while the parietal reach region encodes the target with a predominance of an eye-centered reference frame.

During a visual working memory task, frontoparietal areas can even contain spatial information not related to the body, but relative to a visual object, which is an allocentric form of spatial representation (38). The possibility of such spatial coding in a reach context, which does not reflect the planned physical arm movement, raises the question in which spatial frame of reference frontoparietal cortex represents reach-associated information during BCI learning and whether parietal and frontal areas share a common frame of reference. Rather than reflecting trial-to-trial updates of the corrected movement to produce adapted motor outputs, as observed in the motor and premotor cortices (12,25,39,40), parietal dynamics could reflect the unchanged sensory state of the system during the planning period before movement onset, as part of its role in state estimation (41–47).

Third, we propose that motor adaptation is reflected in changes in motor signals during movement preparation already and that that planning activities co-adapt with signals during movement control. Fast adaptation at the neural level has been previously described from a dynamical system perspective (18,19,48–50). According to this viewpoint, the neural state, which represents the activity pattern across the population of neurons in motor areas, follows established dynamics to generate movements (51–55). During movement preparation, the neural state is initialized with different conditions for different movements and then evolves based on the network-inherent dynamics during movement execution. This concept is particularly applicable to ballistic reaches toward spatially segregated targets in space. The fixed neural dynamics are determined by the stable recurrent connectivity of the network itself (52,56). According to this view, adaptation could be implemented by updating of the initial states, hence, adaptation induced neural changes should reflect in late planning activity.

Over the course of fast adaptation, so the idea, neural dynamics are initialized with changing conditions to compensate for movement errors from trial to trial, while the network structure in the motor cortex remains stable. A BCI learning study in monkeys indicated that such adaptive state initialization probably applies to BCI controlling units in the motor cortex (18). In line with this finding, other work suggested that motor learning involves systematic changes in preparatory activity within the motor cortex to adapt to a force field (57). It is yet unclear to what extent such adaptation of initial states also applies to changes in preparatory activity in the larger frontoparietal network, including non-controlling units and areas outside the motor cortex. By introducing an instructed delay in the BCI movement task, we can compare adaptation of the preparatory states with the movement-associated dynamics. We ask whether such a relationship between the two states exists also for visuomotor adaptation and whether the relationship changes over the course of learning.

In this study, we conducted a 3D visuomotor rotation task under BCI control to investigate neural adaptation across the distributed frontoparietal network, including primary motor cortex (M1), dorsal premotor cortex (PMd), and parietal reach region (PRR). We selectively connected only a subset of units, referred to as controlling units, to the decoder to assess whether neural signatures of adaptation extend beyond the units immediately controlling the BCI movements. After demonstrating visuomotor rotation adaptation in BCI control of 3D movements and its compatibility with the idea of fast within-manifold learning, we address our three main research questions. First, we determine the spatial frame of reference in which the non-controlling units co-adapt with the controlling units that determine the behavior. Second, we test if this co-adaptation also affects the coding in parietal areas, remote from the frontal lobe areas that controlled the BCI movements. Third, we test the hypothesis that re-association learning leads to adaptation of movement planning signals consistent with the observed adaptation in motor behavior.

## Results

### Monkeys learned to control movements in 3D-virtual reality with BCI

To better understand the neural mechanisms underlying movement planning and execution, we designed an experiment to investigate adaptive changes in neural dynamics during a memory-guided reach task under BCI control. In our experiment, two rhesus monkeys (Y and Z) performed a memory-guided 3D center-out reach task using a computer cursor in a virtual reality (VR) environment (Fig. 1A and Supplementary Fig. S1A). The task involved the presentation of a target, followed by a memory period during which the target was no longer visible. The monkeys were then required to move the cursor to the previously cued target in a corner of a 3D cube (Fig. 1B). This delay period between target presentation and movement execution allowed us to investigate adaptive changes in neural dynamics during movement planning, independent of immediate sensory feedback, and compare these to the adapted motor commands during execution.

**Figure 1:**
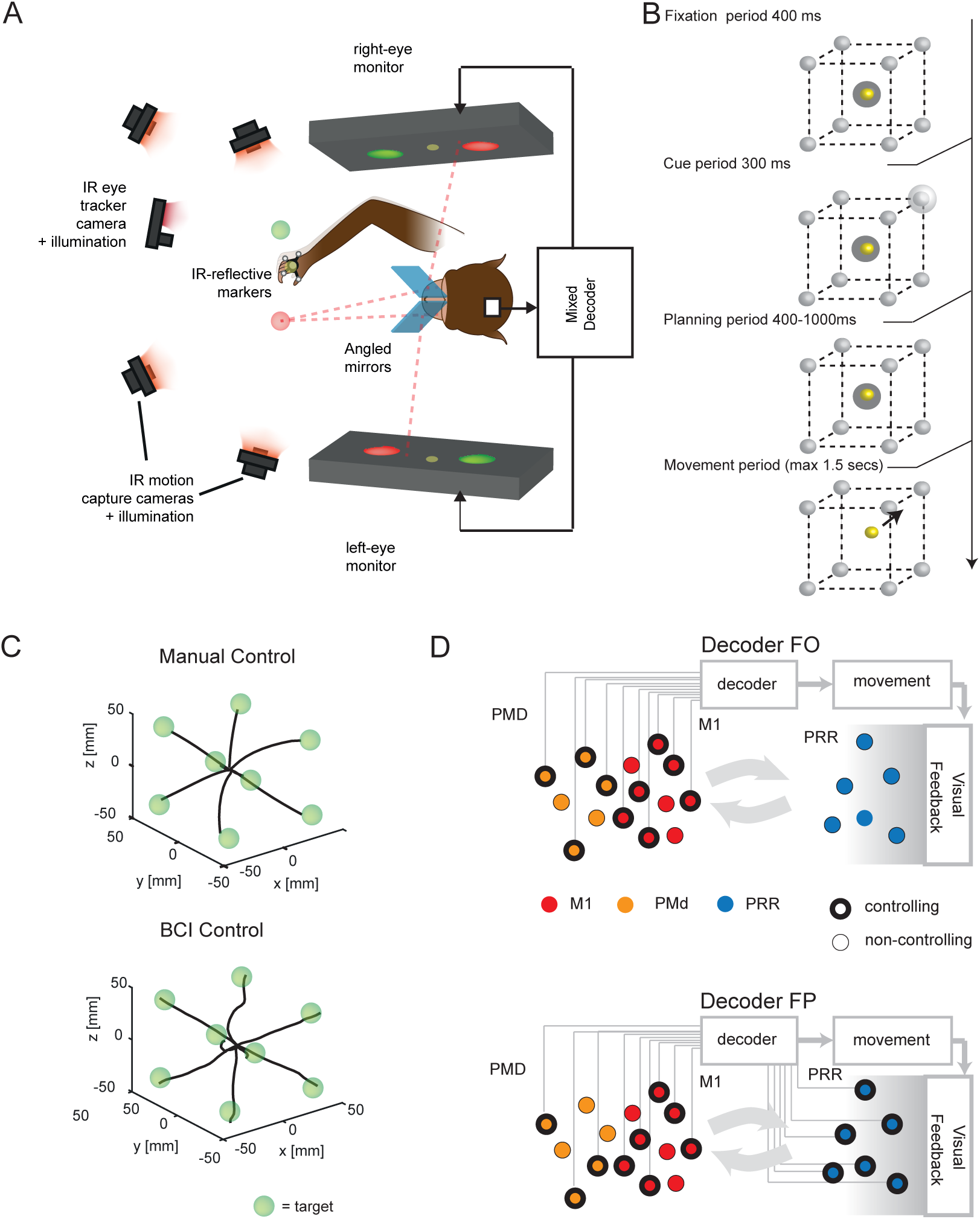
Experimental Setup and Decoding Scheme for 3D Virtual Reality Task. A) Schematic representation of the 3D virtual reality (VR) setup allowing control of movements through manual control (MC) and brain-computer interface (BCI). The setup includes four infrared cameras for online tracking of hand position using reflective markers, enabling realistic 3D movements and decoder calibration for the BCI task. The monkey’s other arm was gently restrained, and gaze position was monitored using an infrared eye tracker. B) Memory-guided center-out reach task in 3D. In each trial, one of the eight corners of the 3D cube was briefly cued as the target and had to be reached after a variable memory period. C) Averaged 3D trajectories obtained from one experimental session in Monkey Y during 3D reaches performed under the brain-computer interface (BCI) condition. D) Two alternative decoding schemes. A subset of M1 and PMd cells was always involved in decoding, represented by black circle outlines. In contrast, PRR cells were either fully integrated into the decoding process (decoder FP) or entirely disconnected (decoder FO).

The monkeys controlled the cursor either through natural hand movements (manual control, MC) or via a BCI driven by intracortical neural activity (Fig. 1C). During MC trials, the cursor movements in virtual space were aligned with the animals’ hand movements in physical space. In BCI trials, the monkeys kept their unrestrained hand in a resting position while controlling the cursor mentally. This ensured a constant somatosensory input during BCI trials. Supplementary Fig. S1B shows that residual hand movements were minimal and barely correlated with cursor speed (Pearson’s correlation: monkey Y, r = 0.085, p < 0.001, n = 575,258; monkey Z, r = 0.072, p < 0.001, n = 108,933). During baseline trials, i.e., without visuomotor perturbation, monkey Y successfully drove the cursor to the target in 99% of the movement trials (trials in which the animal successfully completed the planning period without, e.g., fixation breaks; see Methods) under MC and in 89% of the BCI trials. Monkey Z succeeded in 97% of the movement trials under MC and in 85% of the BCI trials.

We recorded brain activity from three regions: the primary motor cortex (M1), dorsal premotor cortex (PMd), and parietal reach region (PRR). Monkey Y had 64 electrodes in each of M1, PMd, and PRR, while monkey Z had 64 electrodes in M1 and 96 electrodes in each of PMd and PRR (Supplementary Fig. S1C). Units from all three areas were selectively connected (controlling units) or not connected (non-controlling units) to the decoder (Fig. 1D). This design allowed us to investigate whether adaptive mechanisms in individual neurons depended on their immediate causal influence on movement, which is only guaranteed for connected units. Subsets of PMd and M1 units were always involved in the decoder, whereas PRR units were either entirely connected (in combination with PMd-M1; Decoder fronto-parietal (FP), Fig. 1D lower) or completely disconnected (Decoder frontal-only (FO), Fig. 1D upper). This setup enabled us to study adaptive mechanisms in PRR even without its direct involvement in generating movements and to compare it to the case when PRR directly contributes to the control of BCI movements.

### Adaptation of BCI movements was induced by perturbed visual feedback in a 3D visuomotor rotation task

We utilized a short-term visuomotor rotation (VMR) adaptation paradigm tailored to our 3D experimental setup (58) to induce repeated BCI learning (Fig. 2). In the daily sessions, first, the decoder was recalibrated until animals became proficient in controlling unperturbed cursor movements without computer support (see Methods). We then recorded 160 baseline trials before we introduced the perturbation, a rotation of 30 degrees to the visual feedback (cursor) in the fronto-parallel X-Y plane, for 300 trials (Fig. 2A). In each daily session, the perturbation was consistently either clockwise or counterclockwise only (supplementary table, Table S1). The perturbation was applied globally throughout the workspace, which means it affected movements to all eight targets but only in the X and Y dimensions, not in the depth dimension away from the body (Fig. 2A left). Finally, the perturbation was removed during the washout phase, which continued until the animals disengaged from the task (Fig. 2A right).

**Figure 2.**
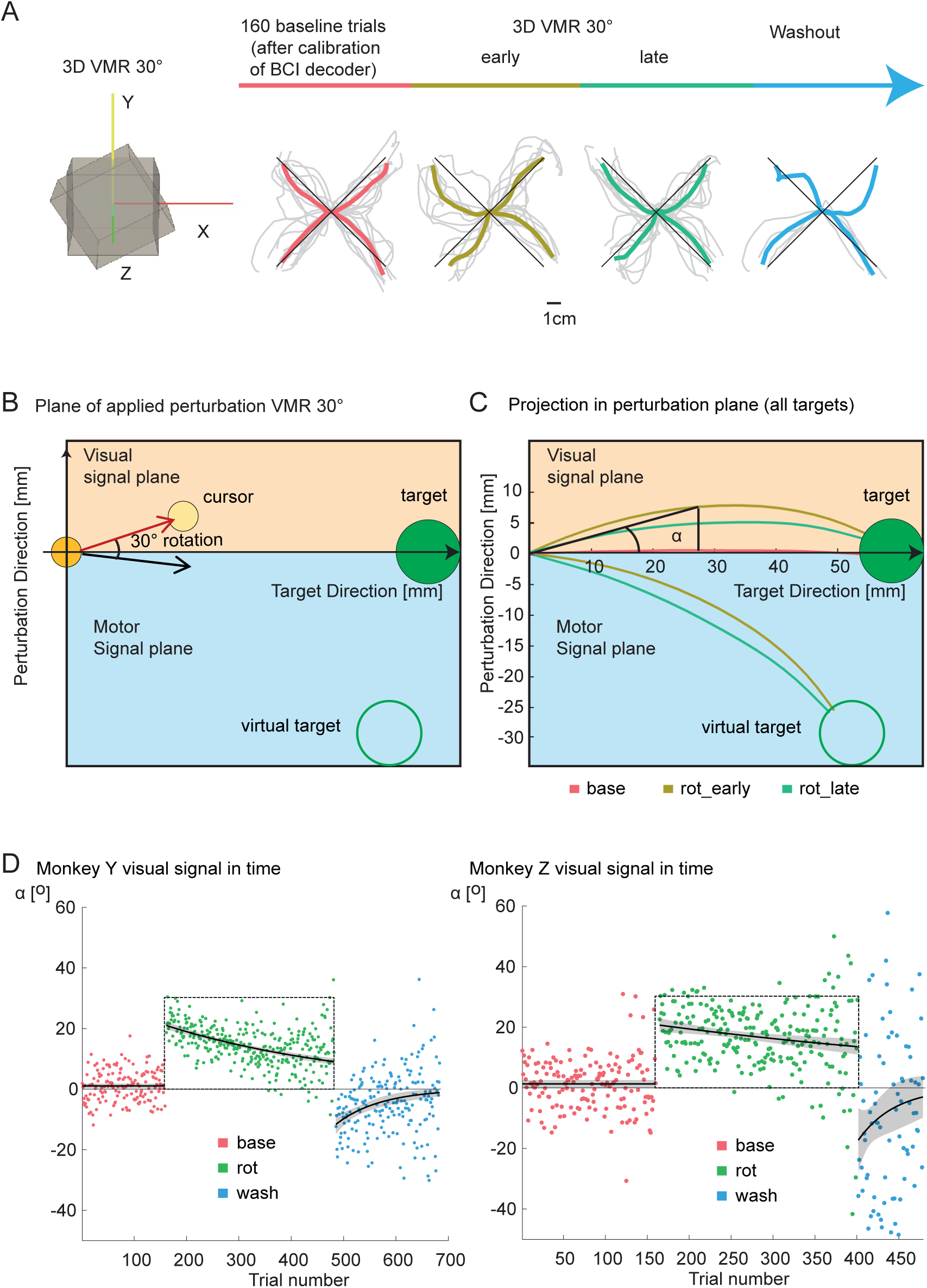
Experimental paradigm and BCI adaptation. A) Left: Visual representation of 3D-VMR. In different experimental sessions, the 3D movements were rotated 30 degrees clockwise or counterclockwise in the frontal-parallel plane. The Z-axis points towards the monkey’s head and was unperturbed. Right: The experimental protocol consisted of a baseline phase followed by a 30-degree VMR and a washout phase where the visual perturbation was removed. Below each phase of the experimental progression, example traces from a single experimental session in Monkey Y are shown. B) In the plane of the applied perturbation, deviations above the zero line correspond to positions in the visual signal plane. Conversely, negative deflections in trajectories exist in the motor signal plane and correspond to the motor commands issued to counteract the visual perturbation. For averaging across all target directions, the trajectories were rotated into a shared reference frame. One axis was aligned with the perturbed direction, while the other was aligned with the vector from the center to the target. In this plane, trajectories above the zero line represent positions in the visual signal plane, while trajectories below zero pertain to the motor signal plane. Additionally, we calculated the angle at the midpoint of each trajectory as the angle between the perturbation direction and the vector from the center to the target. C) For the analysis, the 3D trajectories were projected onto the 2D plane of the applied perturbation. The trajectories showed stereotypical adaptation profiles during rotation and negative after-effects during washout. D) The angle at the midpoint of the trajectory, plotted as a function of trial number, reveals stereotypical adaptation profiles in response to the introduced VMR. The perturbation angle approaches 30 degrees upon perturbation introduction, gradually decreases with learning, and then deviates in the opposite direction during the washout phase following perturbation removal.

To visualize movement adaptation, we projected the 3D movement trajectories of the controlled cursor into the X-Y plane affected by the perturbation (perturbation plane). During the baseline phase, the trajectories were on average relatively straight toward the target. During early rotation trials, as expected as a consequence of the perturbed feedback, trajectories in the perturbation plane curved in the direction of the applied feedback perturbation (Fig. 2A right). The average trajectories were less curved in late rotation trials. In the early washout phase, the trajectories on average curved in the opposite direction compared to during perturbation. To obtain averaged trajectories across targets, we rotated them for each target such that the X-axis aligned with the center-to-target direction and the Y-axis aligned orthogonally to this (perturbation direction, Fig. 2B; see Methods). Figure 2C depicts the average cursor trajectories across all eight targets and for all sessions for monkey Y. The reduced curvature in late rotation trials observed during washout indicate successful adaptation. Additionally, the corresponding “motor” signal, representing the trajectories that would be produced by the unperturbed decoder using the neural activity of the controlling units, is visualized (Fig. 2B–C).

To quantify adaptation, we calculated the trial-by-trial angular movement error (α) from the starting direction of the movement. This was achieved by measuring the angle of the cursor position at the halfway point of the trajectory relative to the straight line connecting the starting position and the target position (Fig. 2C). On average, during baseline trials, the angular error was close to zero and did not decrease over time, as the fitted data with exponential decay did not show significant differences from zero (Monkey Y: intercept = 0.637, p = 0.26, decay = 0.006, p = 0.43, n = 51; Monkey Z: intercept = 1.5, p = 0.26, decay = 0.0029, p = 0.8, n = 15). It increased to about 20 degrees at the onset of the VMR perturbation and gradually decreased as the monkeys adapted their neural activity to regain better control (Fig. 2D). Fitting the data with an exponential decaying function during the perturbation phase indicated significant visuomotor adaptation (Monkey Y: n = 51, intercept = 20.99, p < 0.001, decay = -0.0026, p < 0.001; Monkey Z: n = 15, intercept = 20.69, p < 0.001, decay = -0.0018, p = 0.0017). During the washout phase, movement errors were committed in the opposite direction to the applied perturbation, consistent with a negative aftereffect as typically seen in visuomotor rotation adaptation. Fitting the data with an exponential decay revealed a significant intercept for both animals (Monkey Y: intercept = -11.63, p < 0.001, n = 26; Monkey Z: intercept = -17.255, p = 0.0018, n = 4), indicating significant aftereffects for both animals. These findings suggest a change in the sensorimotor transformation achieved during BCI adaptation trials.

### Preserved covariance structure suggests within-manifold learning during BCI-VMR

In BCI-controlled movements, behavioral changes represented by the cursor on the screen (i.e., the decoder output) directly mirror the activity of the neural population that controls the decoder. When faced with a VMR-type perturbation, this controlling neural group can restore its performance by re-associating certain target directions with different movement directions (13,17). The re-association hypothesis means that the neurons can direct the cursor to the desired target using activity patterns consistent with those observed during the center-out reach task on which they were originally trained, aiming at a direction suited to compensate for the perturbing rotation. In contrast to the re-configuration hypothesis, the covariance structure between different neurons should change very little with re-associations, since the network remains stable. This stability implies that the neural dynamics that define the low-dimensional neural subspace (i.e., the neural manifold), which explains most of the neural variance before adaptation (during baseline), also explain most of the variance during perturbation and after adaptation (washout).

To test the re-association versus re-configuration hypothesis, we calculated the alignment index (55) of the neural manifolds between the baseline trials and the other experimental phases (Supplementary Fig. S2). Alignment should be high in the case of re-association. The alignment index is computed as the ratio of the explained variance when projecting the neural activity from the other experimental phases onto the principal components derived from the baseline activity (see Methods section). We focused on the first four principal components, which capture the majority of the variance in the baseline trials (Supplementary Fig. S3A,C,E,G). Supporting the re-association hypothesis, we find that a substantial portion of the initially explained variance remains constant throughout the learning process. In fact, the average difference between the alignment indices during late rotation and the cross-validated alignment index during baseline is smaller than 1% for all the different neural populations that we calculated (Supplementary Fig. S2). This suggests a preservation of the manifold structure during VMR adaptation in the frontal motor network

### Non-controlling units in M1-PMd encode the same corrective task parameters during adaptation of BCI movement as controlling units

Our primary research goal was to identify the spatial reference frames of movement parameters encoded by neurons without a direct causal link to the movement (non-controlling units) in each brain area. During BCI movements, non-controlling units could encode movement parameters in either a visual coordinate system (reflecting the cursor movement) or a physical coordinate system (reflecting intended limb movements). To distinguish between these spatial encoding strategies, we used an offline decoding approach to reconstruct movement trajectories from non-controlling units.

First, a Kalman filter decoder (as for online decoding) was trained offline with neural activity of the non-controlling units during BCI baseline trials (training set, Fig. 3A) to reproduce the cursor trajectories actually produced by the animal with the controlling units during baseline. During unperturbed baseline trials, these trajectories represent both the motor command (of the controlling units) and visual cursor feedback alike, since both are spatially congruent (identical except for sensory noise).

**Figure 3.**
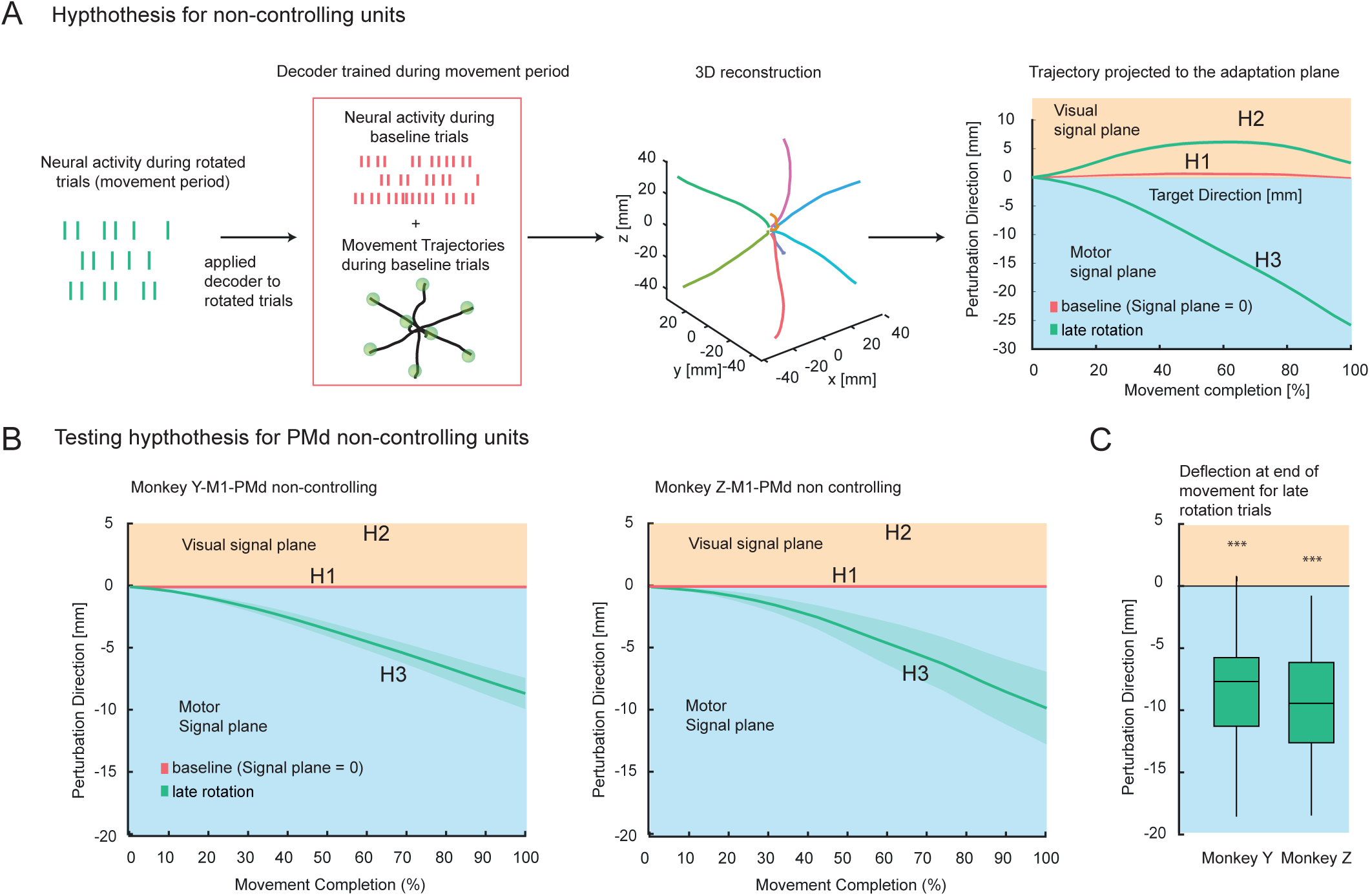
Offline decoder approach. A) Schematic illustration of the offline decoding principle during the movement period. A velocity Kalman filter decoder was applied to reconstruct theoretical memory trajectories from neural activity (first column). During the movement period, real BCI trajectories were used for regression with neural firing rates from non-controlling neurons during baseline trials (second column, training set). After calibration during baseline trials, the decoder was used to reconstruct continuous trajectories during perturbation trials (third column). The example averaged trajectories demonstrate continuous movements toward the target. In the last column, the reconstructed 3D trajectory from the offline decoder is projected onto the perturbation plane for perturbation trials. These projections assess visual-like or motor-like encoding, where positive deviations along this axis correspond to the visual signal, and negative deflections indicate motor output during adaptation. B) Reconstruction and projection of the offline trajectories from M1-PMd in the adaptive dimension for the late adaptation phase. The depicted trajectories represent the averages across all experiments where non-decoding units were collected. The traces display the average (solid color) and the bootstrapped confidence interval (shaded colors) for baseline (pink) and late rotation (last 50% of the trials per session) trajectories (green). C) Box plot showing deflections of the trajectories at the end of movement for both animals during late rotation trials. A significant negative deflection indicates motor-like encoding (* p < 0.05, *** p < 0.001). The whiskers represent the 5th and 95th percentiles.

Second, this decoder was then applied to the neural activity during VMR trials of the same neural population (test set, Fig. 3A). During VMR trials, the motor command of the controlling units and the visual cursor feedback are spatially incongruent. This allows us to test which spatial reference frame the non-controlling units better correspond to. More specifically, the projection of the decoded trajectories along the perturbation axis (deflection measure) allowed us to study whether neural activity of the non-controlling units during adaptation reflects (i) a stationary, non-adaptive signal, (ii) the adapting visual cursor trajectories, or (iii) the adapting motor signal as produced by the controlling units. Straight trajectories to the target as during baseline would reflect a non-adaptive signal in non-controlling populations (corresponding to deviation in perturbation direction = 0 in Fig. 3A). Deviation from baseline deflecting in the positive direction of the perturbation axis would reflect the visual signal (visual signal plane; corresponding to perturbation direction greater than 0), while deflection in the negative direction would reflect the motor command (motor signal space; corresponding to perturbation direction less than 0).

As a result, the negatively deflected reconstructed trajectories in both animals show that non-controlling units in M1 and PMd reflect the motor command similarly to when they directly control the movement via the decoder (Fig. 3B-C). To accurately capture this deflection, we used a trajectory deflection measure in our analysis. This approach differs from the angular movement error (α) we previously calculated to quantify adaptation, where α was measured at the halfway point of the trajectory relative to the straight line connecting the starting and target positions (Fig. 2C). We chose the deflection measure for the reconstructed trajectories because it accounts for the overall shape and direction of the entire movement path.

The mean deflection of the activity of monkey Y at the end of the movement (100%) was -8.69 mm ± 4.43 mm (mean ± standard deviation), significantly different from zero (one-sample t-test, t(50) = -14.010, p < 0.001). For monkey Z, the mean deflection of activity at the end of the movement was also significantly below zero, at -9.88 mm ± 5.27 mm (t(14) = -7.250, p < 0.001).

To gain deeper insight into the level of adaptation achieved by the non-controlling neuronal population, we calculated the angle representing the deviation in the perturbation direction at the final level of adaptation. This angle was measured between the reconstructed trajectory of each population and the direct path to the target. To compare the adaptation levels across different populations, we then scaled the final angle reached by each neural subpopulation relative to that of the controlling unit population and called this the “relative gain.” The relative gain quantifies the deviation along the perturbation axis relative to the level of adaptation achieved. By expressing these deviations as a fraction of the total adaptation level exhibited by the controlling units, we could assess and compare how much each population contributed to adapting to the perturbation. Specifically, the M1-PMd non-controlling population reached a relative gain of 59% for Monkey Y and 94% for Monkey Z, underscoring the significant adaptation observed in the M1-PMd non-controlling population.

### Controlling and non-controlling units in PRR share the same spatial frame of reference during adaptation of BCI movement as controlling units in M1-PMd

We next investigated how VMR adaptation affects PRR activity, focusing on whether PRR’s adaptation-associated neural dynamics share the same spatial frame of reference as the frontal regions. By analyzing the adaptation process in PRR both with direct influence on movement control (decoder FP, controlling PRR units) and without it (decoder FO, non-controlling PRR units), we determined whether PRR predominantly mirrors visual feedback or plays an active role similar to frontal areas. The overall neural yield in PRR was lower than in the combined M1-PMd recordings for both subjects. To still achieve similar baseline performance across conditions and to have enough non-controlling units in PRR for analysis, the controlling PRR neurons were combined with a subset of M1-PMd units in the FP decoder sessions. This experimental strategy allowed us to compare adaptation of PRR while partially causing the movement or not causing it directly at all.

First, we evaluated whether the respective contributions of M1-PMd and PRR to the cursor movement were balanced during FP decoding, or rather predominated by M1-PMd inputs. Each cell’s contribution was determined by multiplying the norm of the vector that translates decoded speed into its firing rate summed with its baseline firing rate (Fig. 4A), a similar approach as described in (Jiang et al., 2020). The distributions of the contribution values differed only modestly between PRR and M1-PMd (Cohen’s d: d = 0.67 for Monkey Y, d = 0.38 for Monkey Z). This suggests that both the PRR and M1-PMd areas similarly drive the motor output of the decoder. We also ensured that after baseline trials, neither of these areas significantly decreased their firing rate during adaptation which might indicate a substantial silencing of the area. Average firing rates between baseline and adaptation did not differ significantly (Mann-Whitney test, p > 0.05; Fig. 4B). Together, these results indicate an active contribution of PRR to decoder adaptation.

**Figure 4.**
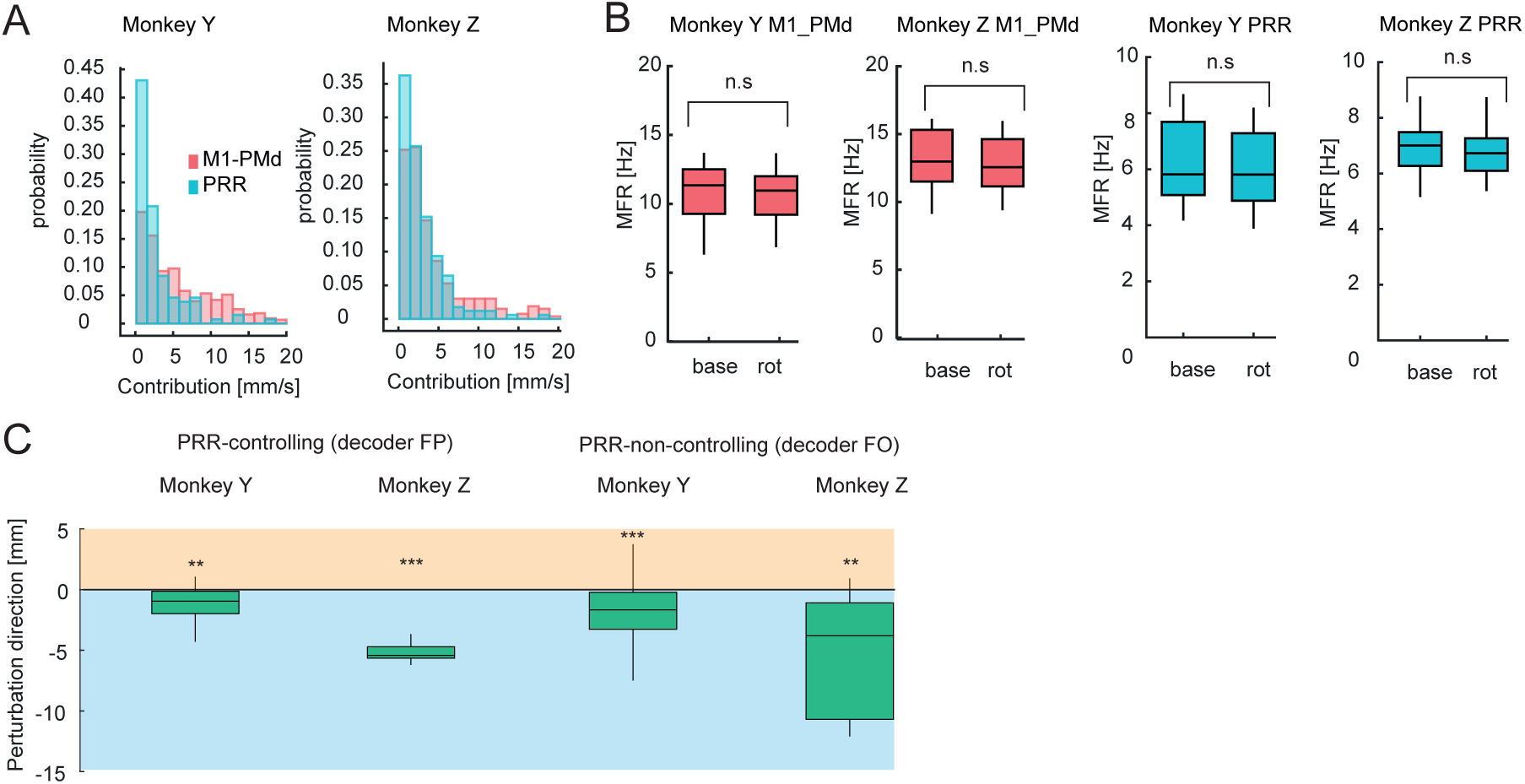
PRR controlling and non-controlling adaptation during movement execution. A) Distributions of the norm of the linear regression vectors from velocity to neural firing rate for every single unit contributing to the decoder, color-coded depending on the neural population. B) Mean firing rates for PRR and M1-PMd controlling units during baseline and late rotation epochs show that neither PRR nor M1-PMd are reducing their firing rate when compared to baseline (all p values non-significant with Mann-Witney test, n = 11 for Monkey Y, n = 5 for Monkey Z). C) Similar to figure 3C, the bar plots show the deflection of the trajectories at the end of the reach. Each distribution consists of all PRR neurons and is tested against zero for the two different decoding schemes and the tow monkeys separately (** p < 0.01, *** p<0.001.)

As a main finding, we see that the changes in PRR in response to the VMR show motor-like characteristics as observed in M1-PMd (Fig. 4C). The mean deflection of the trajectories at the end of the movement under the FP decoder configuration in Monkey Y was -1.64 mm ± 1.56 mm (mean ± SD). A one-sample t-test indicated that this deflection was significantly different from zero (t(10) = -3.48, p = 0.006). For Monkey Z, the mean deflection was -5.15 mm ± 0.932 mm. A subsequent one-sample t-test showed that this deflection was also significantly different from zero (t(4) = -12.36, p = 0.0002). Results were similar when PRR was non-controlling (decoder FO). The mean deflection of the trajectories at the end of the movement for PRR non-controlling units of Monkey Y was -1.85 mm ± 2.25 mm. A one-sample t-test indicated that this deflection was significantly different from zero (t(39) = -5.20, p < 0.001). For Monkey Z, a deflection of -5.16 mm ± 0.93 mm was observed for decoder FO (t(4) = -12.37, p = 0.0001). Additionally, for Monkey Z, the mean deflection for decoder FO was -5.19 mm ± 5.08 mm. A subsequent one-sample t-test showed that this deflection too was significantly different from zero (t(9) = -3.226, p = 0.010). These results indicate a motor-like encoding during adaptation, qualitatively similar to M1-PMd (Fig. 4C). Even when PRR was not contributing to the control of the decoder, an equivalent pattern of adaptation was observed, indicating motor-related changes rather than representation of the visual feedback.

For the PRR controlling units, Monkey Y showed a 13% relative gain to the final adaptation, while Monkey Z showed 64%. When PRR was non-controlling, Monkey Y showed a 14% relative gain, and Monkey Z achieved 37%. In summary, our findings suggest that PRR exhibits substantial motor-like encoding during learning, even when it is not directly controlling the movement, highlighting its involvement in the adaptation process. This indicates that PRR plays a more integral role in motor adaptation than merely reflecting visual feedback.

### Comparing Movement Planning and Execution to Uncover Neural Adaptation Dynamics in BCI Learning

An important aspect of our study design is to compare movement planning and execution to understand the dynamics of neural adaptation. We ask whether motor preparation and the associated initial states of neural dynamics are updated trial-by-trial during adaptation, or if only online motor control changes during BCI learning. First, while the animals were required to maintain ocular fixation during the planning phase of the task, they could still have attempted to move the cursor toward the target in this phase. To rule this out, we verified that the planning phase in the BCI context was comparable to the planning phase in the MC context. We applied a principal component decomposition and cross-projected the neural activity during planning onto the dimensions that best explained the activity during movement. We found that the explained variance in the planning activity, when projected into the first four dimensions of the movement subspace, was significantly reduced in comparison to the movement activity (Supplementary Fig. S3). The subspace misalignment was more pronounced in frontal areas compared to parietal areas in both monkeys. The alignment patterns were equivalent in both manual and BCI trials. These results show substantial differences between the manifold structure in the planning and movement phases not only in MC but also in BCI. This argues against a strategy of monkeys attempting movements during the delay period.

### Motor planning activity in M1-PMd reflects a re-aiming strategy

To better understand how the fronto-parietal network adjusts to altered feedback, we examined changes not just during the control of movement but also during its planning phase. To measure neural changes during planning, we employed an offline decoder approach similar to the one used for studying the movement phase. For planning, similar to movement decoding, we trained the decoder with baseline trials but used neural activity from the memory period. Unlike movement decoding, this training involved regressing the firing rates against a directional vector that continuously pointed to the cued target. Decoding the direction every 50 ms enabled us to reconstruct hypothetical 3D trajectories offline during the planning phase. For each target direction, we took the 400 ms time interval before the ’go’ cue. We then calculated the vector sum of the individual unit vectors, each pointing in the direction decoded within that specific time bin (Fig. 5A). The hypothetical trajectories we produced stem from a direct projection of the neural space into the three-dimensional task space through the decoder, reflecting the intended direction during planning. In essence, these trajectories transform neural activity into a space where variations are explicitly linked to the task space (via the decoder), such as aiming at a target. To validate our decoding methodology during the planning phase, we identified the intended targets by determining the closest target to the final position attained at the conclusion of the planning phase for each reconstructed trajectory within the task space (using an 8-way discrete classifier). In our baseline trials, the decoder consistently produced hypothetical trajectories in the task space that aligned with the intended target (Supplementary Fig. S4).

**Figure 5.**
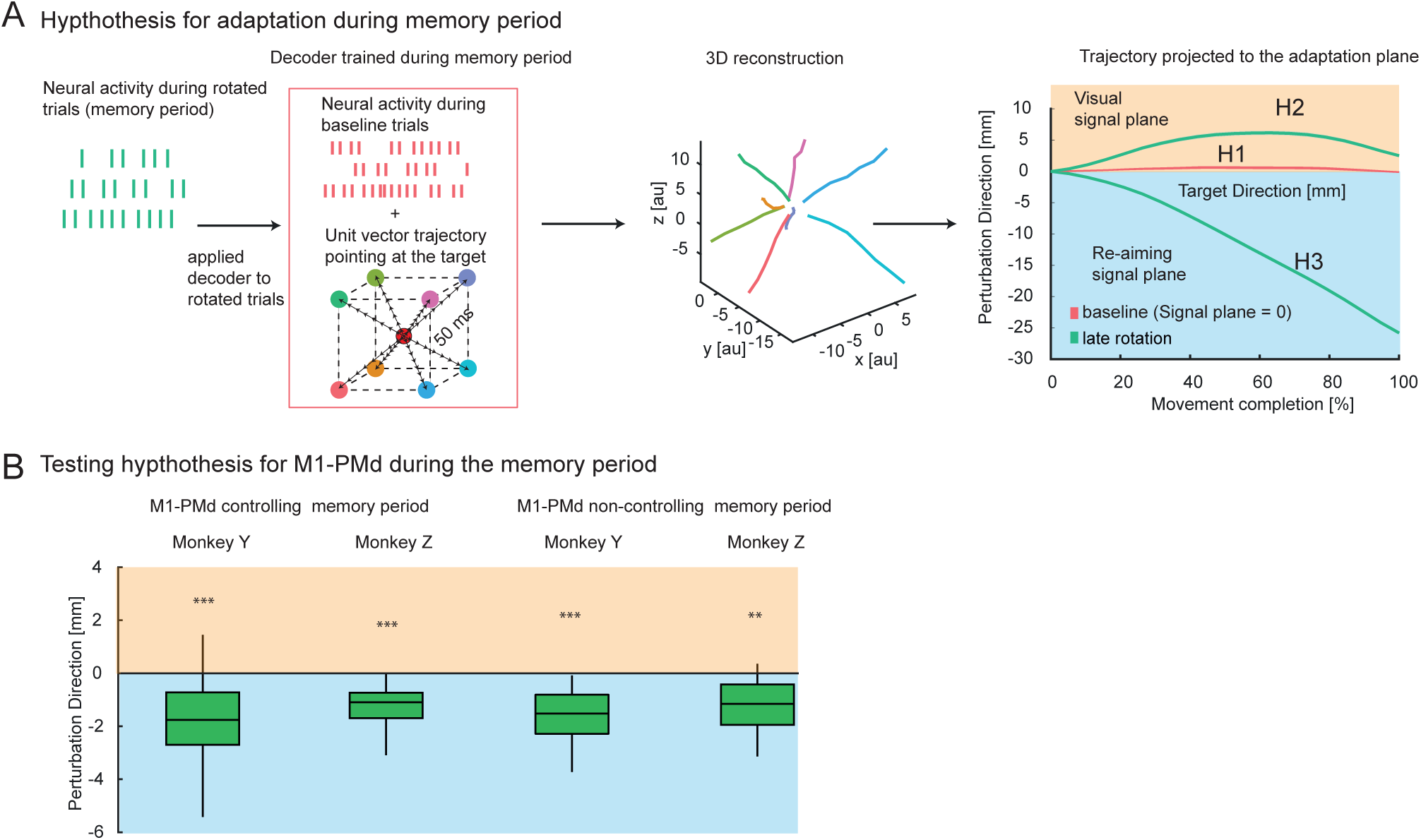
Memory period decoder. A) Schematic illustration of the offline decoding principle during the memory period. As previously described for movement decoding, a velocity Kalman filter decoder was applied to reconstruct theoretical memory trajectories from neural activity (first column). During the memory period, a unity vector always pointing at the target was used for regression with neural firing rates (second column, training set). After calibration during baseline trials, the decoder was used to reconstruct continuous trajectories during perturbation trials (third column). The example averaged trajectories demonstrate continuous movements toward the target. In the last column, the reconstructed 3D trajectory from the offline decoder is projected onto the perturbation plane for perturbation trials. These projections assess visual-like or motor-like encoding, where positive deviations along this axis correspond to the visual signal, and negative deflections indicate motor output during adaptation. B) The reconstructed trajectories in the perturbed dimension exhibit a negative deflection, indicating re-aiming, even during the planning period. At the end of the trajectory, a box plot compares all M1-PMd neurons with zero. The whiskers represent the 5th and 95th percentiles.

To measure the degree of re-aiming due to adaptation, we tested the decoder’s generalization. This was done by training it during the memory phase of the baseline trials and then applying it during the memory phase of the rotation trials. Much like neural activity observed during movement, any deviations in the reconstructed hypothetical trajectories (once projected onto the perturbation direction) would suggest either motor- or visual-like adaptation. We quantified this in a manner equivalent to how we assessed data during movement control (Fig. 5A, right side). For both monkeys, the controlling and non-controlling units in the M1-PMd regions exhibited motor-like adaptation during the planning phase. This suggests the use of a re-aiming strategy (Fig. 5B). In the PRR, there was a general trend towards re-aiming (Supplementary Fig. S5), but this trend was not statistically significant for the individual monkeys.

### Correlated adaptation in planning and movement-related activity strengthens over the course of learning in M1-PMd and PRR

By extracting the extent of re-aiming in both the planning and movement periods, we tested whether the observed level of trial-by-trial re-aiming during planning predicts the re-aiming levels during movement. We correlated the deviations of the decoded trajectories from a direct-to-target trajectory at the conclusion of the planning period with the deviations observed 200 ms after the go cue for each trial (Fig. 6A). We chose the early 200 ms deflection in the movement to reduce the impact of feedback-driven online corrections, which would be anticipated at later intervals. A positive correlation would support the hypothesis that adaptation is mostly achieved by adapting the initial conditions prior to the onset of movement, while movement-associated neural dynamics remain comparably stable. It’s important to note that trajectories in the task space are theoretical during the planning phase and are decoded differently (see Methods) than during the movement phase. Therefore, the regression slope between the planning and the movement trajectories is in arbitrary units. However, relative values in the slope across brain regions, different neuronal groups, or between the initial and advanced stages of adaptation can still signify the ratio at which changes related to adaptation during planning manifest as dynamic changes during movement.

**Figure 6.**
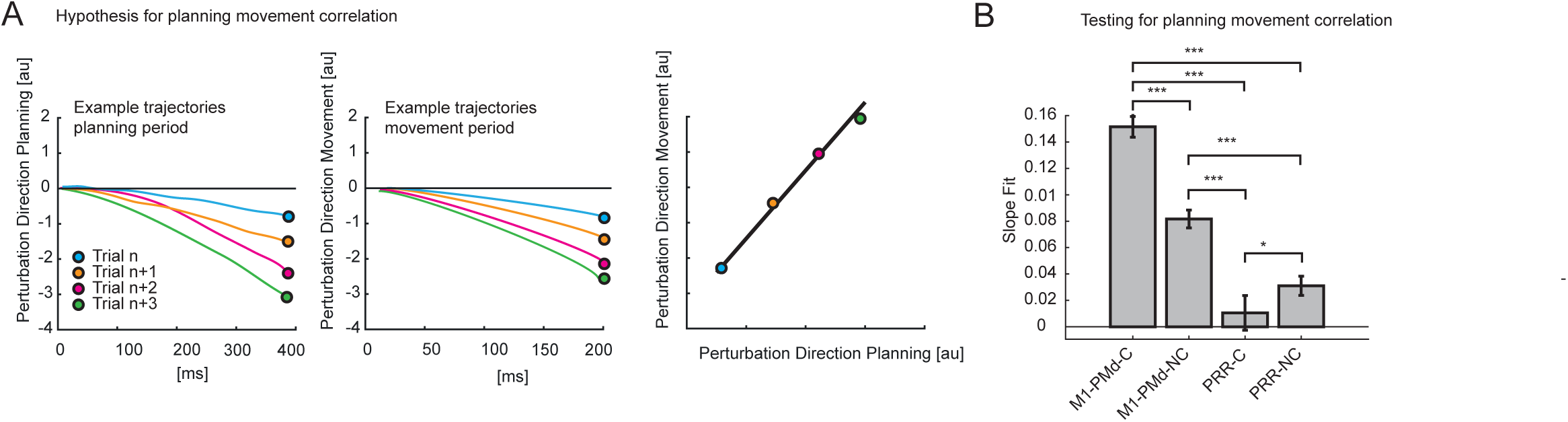
Planning – movement relationship. A. Schematic representation of the analysis method. The reconstructed trajectories at the end of the planning period were regressed against those at 200 ms into the movement phase to test if adaptation levels during planning correlate with those during movement. For each axis, they represent the values assumed for each trial along the perturbation dimension in the specific period (planning and movement). B. Multi-comparison of regression slopes for all controlling and non-controlling populations. The slopes were extracted using a multiple linear regression model from data collected from two animals, with random factors considered. Paired multiple comparisons were deemed significant for an interaction effect between the two tested populations with a Bonferroni-corrected p-value < 0.05 (* p < 0.05, *** p < 0.001). The error bars indicate 95% Bonferroni-corrected bootstrapped confidence intervals as estimated by the model.

We quantified the trial-to-trial relationship between deflections during planning and the direction of subsequent movement using a mixed-effects linear model. This model incorporates the trial number (perturbation trials) as an interaction factor to examine if the relationship evolves during adaptation. We applied a combined model, accounting for both animals, with the individual animal as a random factor and the day of the experiment nested within the animal random factor. Additionally, we separated the analysis for different brain areas and further categorized it based on controlling and non-controlling units. Significant correlations were observed between deflections during planning and movement across all brain areas, but not all subgroups of neurons (Fig. 6B). For controlling and non-controlling M1-PMd populations, slopes were significant (p < 0.001 and p = 0.0085, respectively), indicating correlations. Similarly, for PRR non-controlling units, a significant correlation was observed (p < 0.001). For PRR controlling units, such a relationship was not evident (p = 0.30). When we compared the slope values across all controlling and non-controlling populations in the absence of the interaction parameter, we noticed a significant difference in every comparison (Fig. 6B).

The linear relationship between planning and movement was already apparent during the baseline BCI trials and, though less pronounced, also during the MC trials (Supplementary Fig. S6).

If late planning states and movement are correlated, it could further be that the relationship remains unchanged over time, implying a stable neural coupling that does not adapt with learning. Alternatively, the relationship could evolve as learning progresses, reflecting a dynamic interplay where neural adaptations during planning are linked to changes in movement execution. As learning progresses, the regression slope steepens for both controlling and non-controlling neural populations in PMd-M1, as well as in the controlling population of PRR. This trend is highlighted by a significant interaction with trial number in our model (for M1-PMd controlling p_change_slope < 0.001, for M1-PMd non-controlling p_change_slope = 0.0053, for PRR non-controlling p_change_slope = 0.023). This suggests that, over time, the same degree of re-aiming during the planning phase results in more pronounced motor corrections (data not shown).

## Discussion

Previous studies on motor learning in primates have predominantly focused on the motor cortex in two-dimensional settings (12,18,19,39,49,50). Here we investigated the adaptive mechanisms underlying short-term adaptation in a distributed network involving both frontal and parietal sensorimotor areas. By employing a VMR task in 3D movements, we explored motor adaptation and the associated neural dynamics with the same dimensionality as natural movements (59), avoiding physical constraints. To minimize potential confounds arising from changes in cross-modal congruency due to the applied perturbation, we employed a brain-computer interface (BCI) paradigm which removes movement-contingent proprioceptive feedback. By selectively manipulating the remaining task-relevant visual feedback and exclusively controlling movements through observed neurons feeding into the BCI decoder, we gained precise control over the sensorimotor loop and associated transformations (15,16). Furthermore, we utilized memory-guided reaches to examine how motor planning activity and its relationship to movement are influenced by learning in both frontal and parietal sensorimotor regions. Our findings revealed coherent neural adaptation in a motor-reference frame across both frontal and parietal motor areas. This means, the adaptive mechanism extended to the parietal cortex, a region remote from the frontal neurons controlling the BCI. This adaptation was observed in both planning- and movement-related activity, regardless of whether the neurons controlled BCI movements or not. Our results suggest the presence of a distributed mechanism of “re-aiming” within a motor-reference frame, influencing movement-planning activities even in the parietal cortex.

### Preservation of correlated frontoparietal network structure during VMR adaptation

A key finding of our study on BCI-VMR is that even neurons in the parietal reach-related area (PRR) exhibit changes in neural activity that reflect the modified motor command during adaptation, rather than the changing sensory feedback. Notably, this change is evident even though PRR neurons do not directly govern BCI movements and are remote from the controlling units in PMd-M1. By maintaining a consistent physical state (posture) during BCI movements, we ensured that proprioceptive input remained constant. Consequently, sensory changes induced by perturbations were confined to the visual cursor, which was directly and solely managed by the controlling units via the decoder (16). Therefore, the decoder input represents the motor command, while its output corresponds to the task-relevant sensory feedback. Additionally, using offline decoding, we could analyze the encoding in groups of neurons not involved in online control.

Earlier research has highlighted the preservation of covariance and neural manifold structures during short-term BCI learning tasks in the motor cortex’s controlling units (14,17). We expand this notion by demonstrating that both non-controlling motor cortex units and PRR units adapt, but, crucially, within a consistent correlation structure (Supplementary Fig. S3). This within-manifold adaptation suggests that functional connectivity remains relatively unchanged during VMR learning (14,60,61). Our findings indicate that a correlated network structure is maintained within the widespread frontoparietal sensorimotor cortices during fast BCI adaptation and that this phenomenon is not restricted to network nodes with direct connections to the decoder.

### Consistent encoding of corrective variables in controlling and non-controlling motor units during adaptation

Our findings further indicate that non-controlling M1-PMd and PRR units reflect adaptive changes in the same motor-related spatial frame of reference that controlling units use; in other words, the changes in non-controlling units predominantly reflect the adapted movement directions that the controlling units have to produce to counteract the feedback perturbation. The uniform encoding of the movement-associated corrective variable across the frontoparietal network may facilitate efficient information exchange within a closed-loop system for motor corrections, especially involving direct PRR engagement (62). Our results add to previous work in which parietal area 5 was found to encode motor errors, and where electrically stimulating this area generated movements opposite to the preferred direction of the stimulated cells (63), in line with results from motor areas M1 and PMd (64). The conclusion that PRR generates a signal in the same frame of reference as the motor areas, in our case, is based on the analysis of neural population responses during a motor adaptation task, without a potentially confounding co-varying proprioceptive feedback.

In rule-based tasks that are likely prompting explicit learning of stimulus-response associations, namely an anti-reach task (65), a neural mechanism has been shown in BCI experiments in monkeys (13) and humans (20) that suggests a re-aiming strategy similar to our observations. These previous BCI studies used a discrete classifier to identify which of two possible targets is selected, i.e., they did not provide movement-contingent feedback (the “cursor” was only shown at the beginning and end of the “movement”). The authors refer to the adaptive mechanism as “intrinsic variable learning”, which is contained within a preexisting neural structure and utilizes the preexisting repertoire of neural dynamics. In contrast, our study employed a VMR adaptation paradigm, which typically induces implicit learning of novel motor mappings (29,58,66,67). This suggests that re-aiming-like neural adaptation mechanisms could serve both implicit and explicit motor learning.

### Re-aiming Signals during Planning and Movement Control

In our study, we probed the trial-by-trial adaptation within the frontoparietal network during the movement planning phases. Specifically, we were keen to understand how re-aiming processes during planning are transferred to the execution of movement. Drawing from dynamical systems theory, we considered the possibility that neural activity during planning could establish the initial conditions, thereby guiding the dynamical system to produce adapted movements without the need for explicit cognitive computation (53) or enhanced restructuring of the network.

Our observations revealed linear correlations between trial-by-trial adjustments of planning and movement activity, evident in both controlling and non-controlling units of frontal motor areas M1-PMd and, less pronounced, in non-controlling units of PRR. From a dynamical systems perspective this suggests that the adaptation of initial states to define updated starting conditions for the following dynamic evolution of states during movement might be particularly true for frontal, but less for parietal areas. Additionally, we found an enhancement of this relationship over the learning phase, indicating that changes during planning translated to more pronounced changes in movement in the later adaptation phase. This suggests that not only the motor error was reduced but also that the adaptive mechanism became more efficient over the course of learning.

These findings partly challenge the notion of a linear dynamical system in motor learning, suggesting that the dynamical system itself may undergo changes. However, Kao et al.’s recent theory (68) offer an complementary perspective, proposing that evolving relationships may reflect the brain’s effort to optimize its energetic costs. In this model, efficient control strategies favor selectively targeting dimensions that most impact future motor outcomes, reducing redundant preparatory effort. Supporting this hypothesis, we observed that as learning progressed, adjustments in planning became more effective in driving corrective changes in movement. This shift implies that less extensive re-aiming during planning could still enable more substantial corrections during movement, aligning with the view that learning serves not only to reduce error but also to refine the efficiency of neural control mechanisms in an energetically optimal way.We conducted a comparative analysis of online and offline decoding, revealing that all neurons, including non-controlling units, coherently encode a corrective variable during motor control to counteract visuomotor rotations. This re-aiming or re-association mechanism, previously observed in motor areas, supports efficient information transfer in a feedback control-loop (12,13,17,20,25). Additionally, our results indicate that re-aiming during planning can predict movement correction, although their relationship varies throughout learning and depends on the area and causal relationship between the area and the BCI movement. During the later stages of adaptation, it appears that less re-aiming is required during planning to achieve a specific degree of movement adaptation.

In conclusion, our study demonstrates that generalized adaptation mechanisms operate within a distributed neural network during three-dimensional movements, offering insights into neural manifold structures and motor error correction. By employing a visuomotor rotation (VMR) task in a 3D workspace and using a brain-computer interface (BCI), we captured spatial encoding and motor adaptation without physically constraining movements and independently controlling sensorimotor transformations from proprioceptive feedback.

Our findings highlight the critical role of an integrated motor and parietal sensorimotor network in adaptation. Both frontal and parietal regions primarily encode adapted motor commands rather than merely responding to perturbed visual feedback, supporting the re-association hypothesis of VMR learning. Moreover, learning-associated changes occurred not only in neurons controlling the BCI output but also in neurons not directly connected to the BCI, indicating a widespread mechanism of motor adaptation.

Finally, neural adaptation manifests in both planning and movement-related activities across the brain, with learning-associated changes in the correlation between these phases. This relationship evolved particularly within frontal areas during learning, suggesting different adaptation trajectories in frontal versus parietal regions. Our results underscore the distributed nature of adaptation within neural networks underlying motor control and highlight the importance of this network in fine-tuning spatial encoding for precise movement correction.

## .Materials and Methods

### Stereoscopic 3-dimensional virtual reality setup

The setup allowed the animals to perform reaching movements in a three-dimensional virtual reality environment (58,69). The animals were seated in a dark room with two monitors (BenQ XL2720T, 27-inch diagonal, 1920x1080 px, 60 Hz refresh rate) positioned on either side (Fig. 1A). They looked through two semi-transparent mirrors (75 x 75 mm, 70R/30T, Edmund Optics) angled at 45° relative to the monitors. This created a stereoscopic three-dimensional virtual impression in front of the subjects. The monitors were tilted 30° relative to the horizontal plane so that the workspace was projected below eye level, allowing the animals to perform ergonomic arm movements in front of their body.

To optimize the perception of the virtual workspace, the interpupillary distance of the monkeys was measured (Monkey Y = 33.2 mm, Monkey Z = 33.1 mm). The setup software and screen projections were then calibrated to these individual values to minimize discrepancies between the images for the left and right eyes. Eye position was tracked at 2 kHz (EyeLink 1000 Plus, SR Research LTD, Ottawa, Canada), and the gaze was required to be maintained on the central fixation point during the memory period (see below).

### Manual and BCI control software

The custom-written task controller was implemented in C++, allowing simultaneous tracking of multiple interfaces including the brain-computer interface (BCI), the eye and the hand tracking systems (70). The BCI was implemented as a Matlab program running an online loop, which communicated with the task controller via a Matlab implementation of the VRPN library through a MEX file (71). Neural data were not just recorded to hard drives but also processed online through a MEX interface provided by the recording system (Cerebus, Blackrock Microsystems, Salt Lake City, USA) to extract spike counts (see below). The loop operated every 50 ms in consecutive non-overlapping time windows. During BCI trials, the cursor was automatically placed at the central fixation point during the holding phase and after the movement was completed (see task description below).

### Tracking system calibration

Four infrared cameras tracked the position of four passive markers centered on the monkey’s palm in real-time (Fig. 1A) at 100Hz. The two animals were trained to wear a 3D-printed plastic framework containing the four reflective markers arranged around the right wrist. The center of geometry of the four markers was tracked using four Vicon Bonita B10 cameras (Vicon, Oxford, UK) and aligned with the midpoint of the palm. To facilitate accurate tracking and control in the virtual reality setup, we performed two calibrations. First, we calibrated the cameras to stream the center of geometry of the markers attached to the monkey’s wrist. We added an extra marker at the position of the monkey’s wrist, which served as the reference point during this calibration. Using Vicon Tracker 1.3.1 software (Vicon), we adjusted the center of geometry of the four markers to align with this additional marker. This allowed the software to stream the position of the center of geometry online to the task controller, ensuring that this position corresponded accurately to the monkey’s wrist position in the physical setup.

Second, we calibrated the camera space with the task controller space to ensure proper alignment between the virtual environment and the physical setup. During this calibration, a human operator co-registered the cursor position and the displayed target positions using semi-transparent mirrors. This process translated the coordinates from Vicon space to the task space, allowing the virtual reality system to accurately reflect the monkey’s movements within the task environment.(58).

### Center-out reach task

Two Rhesus monkeys performed a three-dimensional, memory-guided, center-out reach task (Fig. 1B). At the beginning of each trial, during the decoder calibration phase, the animals were required to acquire and maintain a central fixation point with both their hand and gaze for 400 ms (fixation period). Following this, one of eight targets, positioned at the vertices of a 70 mm-sided cube and centered around the fixation point, was briefly displayed for 300 ms (cue period). The animals had to continue holding their hand and gaze at the center of the cube for an additional 400 ms (planning period), followed by a variable delay of up to 600 ms. After the delay, the central fixation target disappeared, signaling the animals to move toward the target (movement period), with a timeout limit of 1500 ms.

During brain-computer interface (BCI) control trials, the cursor movement was controlled by a neural decoder. The task remained identical to the manual control task that had been used for calibration. If, at any point, the animals directed their gaze outside a 30 mm tolerance window around the fixation point or moved their hand more than 15 mm from its starting position (security radius) during the fixation period, the trial was aborted. Manual control trials were employed to calibrate a biomimetic velocity Kalman filter (KF) decoder, which was subsequently used to perform the BCI trials (Fig. 1C, BCI).

Once the animals became proficient in controlling the cursor during the BCI task (reaching a baseline of 160 successful trials), we introduced a visuomotor rotation (VMR) to perturb the visual cursor feedback. This was implemented by applying a 30° visuomotor rotation of the cursor’s movement in the fronto-parallel plane. The direction of the rotation (clockwise or counterclockwise) varied between days, and the perturbation lasted for a variable number of trials with a maximum of 320 trials. Following the perturbation phase, a washout phase was introduced, during which the visuomotor rotation was removed, and the animals returned to controlling the cursor without any rotation for another variable number of trials until they finished their session for the day.

### Neural recordings

Neuronal activity was recorded from two macaque monkeys implanted with floating microelectrode arrays (FMAs, MicroProbes, Gaithersburg, USA) in three cortical areas: M1, PMd, and PRR. Monkey Y was implanted with two 32-channel FMAs in each area, while monkey Z was implanted with three 32-channel FMAs in PMd and PRR (see Supplementary Fig. S1C). Both animals were implanted in the left hemisphere, contralateral to the arm used for calibrating the BCI decoder and performing the manual control task (Supplementary Fig. S1B).

In single-day sessions, we recorded from 128 electrodes simultaneously in monkey Y, while we recorded from all 256 electrodes in monkey Z. Activity was recorded online and stored on a hard drive using a 128-channel Cerebus system (Blackrock Microsystems, Salt Lake City, USA) for monkey Y and two 128-channel Cerebus systems for monkey Z. Data were sampled at 30 kHz. Signals were band-pass filtered (250 Hz - 7.5 kHz), and a threshold of -4.5 times the root-mean-square (RMS) voltage was used to isolate below-threshold waveforms.

Before the start of each daily experimental session, manual online sorting was performed by an expert user. Sorting involved identification of clusters in principal component (PC) space to classify neuronal activity based on waveform characteristics.

### Neural decoder

Online neuronal activity during movement was used to train a velocity Kalman Filter (KF) decoder (72) by regressing neuronal firing rates from a subset of single and multi-units with hand velocity (50 ms steps, non-overlapping time windows) while the monkey manually performed the task during the movement period. During FO decoding a subset of M1 and PMd units where used while during FP decoding subsets of M1 and PMd units were combined with all PRR units. After an initial manual control (MC) calibration consisting of about 60 trials for initializing Kalman filter parameters, the velocity KF decoder was retrained using velocity vectors that directly pointed toward the target (73). During retraining, first the computer mostly controlled the cursor movement. The computer’s contribution then was step-by-step progressively reduced until real-time neural activity fully guided the cursor. The transition was managed by applying a weighted vectorial sum of both the computer-generated and neural signals, gradually increasing the brain’s control for smoother cursor operation. This adjustment was performed over successive blocks of 20–30 trials, reducing the computer’s contribution from 70% to 0%. The process was not automated and required expert judgment to determine the appropriate timing of the transitions. Additionally, hand movements greater than 15 mm from the resting position were discouraged to prevent the animal from using its hand during BCI trials. This retraining calibration step was necessary to eliminate hand movement during BCI control.

### Projection of trajectories on the adaptive space

We employed the method described by Jarosiewicz et al. (25) to average trajectories reaching different targets. In brief, trajectories were first projected onto the plane of the applied perturbation (xy – fronto-parallel plane). For each trajectory to a target, the dimension orthogonal to the direction from the center to the specific target was defined as the “perturbation dimension.” This allowed all trajectories to the eight different targets to be averaged within this common reference frame. The positive sign of this second axis was chosen based on the direction of the applied perturbation (clockwise or counterclockwise, depending on the session).

### Neural data pre-processing (offline)

The online-sorted spikes were re-sorted offline using Boss software (v.1.0.3, Blackrock Microsystems, Salt Lake City, USA) to better isolate and identify additional units among the non-controlling units. Units used for online decoding were left unchanged. A threshold of -4.5 times the standard deviation of the noise was used to isolate spikes, with a refractory period of 1.5 ms. Clusters were identified using a semi-automated k-means method in principal component (PC) space. This semi-automated process required the user to specify the number of clusters and define the initial centroid for each.

### Offline trajectory reconstruction

To study changes in the encoded correction variable during adaptation for non-controlling units, and during the memory period for all units, we applied a Kalman Filter (KF) decoder offline, similar to the one used for online BCI trials, to reconstruct cursor trajectories. The decoder was always trained on the activity from baseline trials and then applied to the activity from the rotated trials to identify the encoded variable during adaptation.

For the evaluation of the decoder during baseline trials, a leave-one-out cross-validation procedure was used to include all baseline trials in the analysis. Each time point, sampled every 50 ms and containing all the spikes from each neuron, was used as input to the decoder. For the decoder’s training output, real movement speeds were used during the movement epoch (sampled every 50 ms), while a normalized vector pointing to the cued target was used during the planning epochs (also sampled every 50 ms). This approach allowed us to reconstruct theoretical neural trajectories from the decoded speeds during movement for non-controlling units and during the memory phase for both controlling and non-controlling units.

### Manifold and alignment index

We calculated the principal component space for each analyzed epoch and region, plotting the variance explained by each dimension (Supplementary Fig. S3), averaged across the dataset. In the cross-projected analysis of Supplementary Fig. S3A, C, E, G, we computed the principal components (PCs) during the movement phase, projected the activity from the memory phase onto the seven principal components, and calculated the explained variance in each specific direction (component). For the alignment index calculation in Supplementary Fig. S3B, D, F, H, we followed the method of Elsayed et al. (2016), considering the top four PCs in the transformation.

In brief, the alignment index measures manifold alignment and is calculated as the ratio between the explained variance of the projected activity (e.g., during the planning phase) along the original dimensions and the explained variance of the original main dimensions (e.g., derived from movement activity). To calculate the original principal component space, we used cross-validated baseline trials to ensure that the alignment index reflects genuine differences between baseline and experimental phases, avoiding artifacts from overfitting.

### Decoder contribution

From the matrix estimated in the online Kalman filter decoder, which relates cursor movements to firing rates, we determined the firing rate of a single cell given the direction of movement using the following equation:

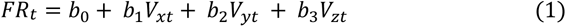

We calculated the amplitude of the regressing vector (b1,b2,b3),summed to b0, to estimate the single neuron’s contribution to the decoder, as shown in Figure 5A.

### Generalized linear model

We used a linear model to test the significance of the relationship between the planning and movement phases in brain each area, separately for the groups of controlling and non-controlling units. We also included trial number as variable to model slope changes due to learning and tested the interaction between learning and the planning-movement slope. Different experimental sessions and the two animals were treated as random factors in the model, with a nested hierarchical structure. The Wilkinson notation for the modelled variable was as follows:

Re-aiming_movement ∼ 1+ Re-aiming-memory + Re-aiming-memory:trial_number + (1|animal/ recording_session)

To calculate slope differences between controlling and non-controlling populations, as well as between different areas, we employed a similar model that incorporated pairwise-tested interaction effects for the areas being compared. All six pairwise comparisons were corrected for significance using the Bonferroni method (see Figure 5B). The corresponding Wilkinson notation was as follows:

Re-aiming_movement ∼ 1 + Re-aiming-memory + Re-aiming-memory:area + (1|animal/ recording_session)

## Acknowledgements

German Research Foundation (DFG, Germany, Grant Number FOR-1847-GA1475-B2), received by AG. German Research Foundation (DFG, Germany, Grant Number SFB-889), received by AG. Federal Ministry for Education and Research (BMBF, Germany, Grant Number 01GQ1005C), received by AG. Federal Ministry for Education and Research (BMBF, Germany, Grant Numbers 01GQ0814), received by AG.

## Author contributions

E.F., A.G. and P.M. conceived the study. E.F. analyzed data and prepared the figures. E.F, A.G. and P.M. wrote the manuscript. E.F and P.M built the 3D-BCI setup. All authors reviewed the manuscript.

## Competing interests

The authors declare no competing interests.

## Data availability

The datasets used and/or analyzed during the current study are available from the corresponding author on reasonable request.

## Notes

### Competing Interest Statement

The authors have declared no competing interest.

### Summary of Updates

corrected figure position since the text in the bioRxiv template overlapped with the figures

