## Supplementary information for "Planning and movement activities co-adapt in a motor-reference frame during 3D BCI-controlled reach adaptation in monkey frontal and parietal cortices"

**Supplementary table T1. Experimental design.**

Table T1 summarizes, for each recording session, the number of units for each population, the type of decoder used, the rotation angle, and the direction of rotation (clockwise, CW, or counterclockwise, CCW).

**Supplementary Figure S1. Hand Movement Speed and Array Implantations**

A) VR setup used for concurrent manual control (MC) and brain-computer interface (BCI) trials.

B) Comparison of real hand movement speed during BCI trials (cyan) and cursor speed during BCI trials (pink). The graph displays the speed of hand movements executed by the monkey during BCI trials as well as the cursor speed during BCI trials, along with the Pearson correlation coefficient indicating the very low correlation between these two speeds.

C) Array locations. Monkey Y received six 32-channel floating micro-wire arrays (FMA, MicroProbes for Life Sciences, Inc., USA) in three distinct brain regions contralateral to the arm used for decoder calibration.

Two arrays with staggered electrode lengths ranging from 7.1 to 1.9 mm were implanted in the arm region of the primary motor cortex (M1), two with staggered electrode lengths ranging from 4.5 to 2.1 mm in the dorsal premotor cortex (PMd), and two with staggered electrode lengths ranging from 7.1 to 1.9 mm in the parietal reach region (PRR). Similarly, in Monkey Z, three arrays were implanted in PMd, two in M1, and three in PRR.

**Supplementary Figure S2. Alignment index during task epochs**

Alignment indexes (AI) during various task epochs for planning (left) and movement (right) phases were calculated. The AI was determined by projecting the neural activity from both the early and late adaptation phases onto the principal components (PCs) derived from baseline activity. We selected a sufficient number of PCs to explain 90% of the baseline variance. AI during the baseline was computed using cross-validation. Notably, AI exhibited minimal variation between the baseline and visuomotor rotation (VMR) phases, indicating the preservation of neural manifolds throughout the learning process.

**Supplementary Figure S3. Neural manifold comparison between MC and BCI**

A) Explained Variance: Monkey Y's neural activity during MC planning was projected onto principal components calculated from neural activity during movement. The explained variance for both activities used to calculate the components (movement training, green) and for cross-validated trials not used in component calculation (movement test, blue) is shown. Bars show 95% confidence intervals.

B) Alignment Index (Monkey Y - MC): Alignment index for Monkey Y calculated during MC for M1-PMd (left) and PRR (right). It measures the extent of overlap among neural manifolds during planning and movement execution using the first four principal components. The whiskers represent the 5th and 95th percentiles.

C) Explained Variance (Monkey Y - BCI): Similar to A, but for BCI trials.

D) Alignment Index (Monkey Y - BCI): Similar to B, but for BCI trials.

E) Explained Variance (Monkey Z): Same analysis as A, but for Monkey Z.

F) Alignment Index (Monkey Z): Same analysis as B, but for Monkey Z.

G) Explained Variance (Monkey Z - BCI): Same as C, but for Monkey Z.

H) Alignment Index (Monkey Z - BCI): Same as D, but for Monkey Z.

Supplementary Figure S4: Memory period decoder

A) For Monkey Y, averaged trajectories of the last 50% of trials reconstructed with the memory period decoder, similar to Figure 5, but for PRR controlling and non-controlling units.

B) Same as in A), but for Monkey Z.

**Supplementary Figure S4: Memory period decoder**

A) For Monkey Y, averaged trajectories of the last 50% of trials reconstructed with the memory period decoder, similar to Figure 5, but for PRR controlling and non-controlling units.

B) Same as in A), but for Monkey Z.

**Supplementary Figure S5-Target classifier from memory period decoder**

Using the method described in Figure 5, we accurately classified the cued target with performance significantly above chance level (12.5%, corresponding to one of eight possible options). This classification was achieved by analyzing the final position of the trajectory within one of the height-specific cubes at the end of the memory period.

**Supplementary Figure S6. Planning – movement relation during baseline and manual control**

Similar to the planning-movement relationship during adaptation, a linear relationship between the two phases can already be observed during manual control (MC) trials and BCI trials.

| Date | M1_PMD_C | M1_PMD_NC | PRR_C | PRR_NC | Rotation_angle | Monkey | Decoder | Rotation_type |
|---|---|---|---|---|---|---|---|---|
| 2016-07-06 | 44 | 12 | | 37 | 30 | Y | FO | CCW |
| 2016-07-08 | 38 | 13 | | 30 | 30 | Y | FO | CCW |
| 2016-07-27 | 45 | 6 | | 48 | 30 | Y | FO | CCW |
| 2016-07-28 | 48 | 16 | | 51 | 30 | Y | FO | CCW |
| 2016-08-05 | 40 | 24 | | 39 | 30 | Y | FO | CCW |
| 2016-08-09 | 42 | 26 | | 38 | 30 | Y | FO | CCW |
| 2016-08-10 | 36 | 25 | | 32 | 30 | Y | FO | CCW |
| 2016-08-13 | 30 | 27 | | 42 | 30 | Y | FO | CCW |
| 2016-10-07 | 43 | 10 | | 28 | 30 | Y | FO | CCW |
| 2016-10-12 | 41 | 23 | | 28 | -30 | Y | FO | CW |
| 2016-10-27 | 42 | 8 | | 23 | -30 | Y | FO | CW |
| 2016-11-02 | 38 | 13 | | 18 | -30 | Y | FO | CW |
| 2016-11-03 | 35 | 18 | | 23 | -30 | Y | FO | CW |
| 2016-11-04 | 30 | 20 | | 30 | 30 | Y | FO | CCW |
| 2016-11-09 | 30 | 16 | | 22 | 30 | Y | FO | CCW |
| 2016-12-07 | 43 | 12 | | 13 | 30 | Y | FO | CCW |
| 2016-12-09 | 41 | 19 | | 19 | 30 | Y | FO | CCW |
| 2016-12-20 | 48 | 12 | | 15 | -30 | Y | FO | CW |
| 2016-12-21 | 38 | 14 | | 20 | 30 | Y | FO | CCW |
| 2017-01-04 | 44 | 16 | | 13 | -30 | Y | FO | CW |
| 2017-01-05 | 51 | 12 | | 12 | 30 | Y | FO | CCW |
| 2017-01-11 | 43 | 19 | | 14 | -30 | Y | FO | CW |
| 2017-01-15 | 49 | 18 | | 15 | 30 | Y | FO | CCW |
| 2017-01-18 | 49 | 17 | | 14 | -30 | Y | FO | CW |
| 2017-02-23 | 42 | 25 | | 12 | 30 | Y | FO | CCW |
| 2017-02-24 | 43 | 25 | | 10 | 30 | Y | FO | CCW |
| 2017-03-01 | 31 | 27 | | 8 | -30 | Y | FO | CW |
| 2017-03-14 | 46 | 18 | | 8 | -30 | Y | FO | CW |
| 2017-03-16 | 38 | 16 | | 10 | 30 | Y | FO | CCW |
| 2017-03-29 | 38 | 12 | | 8 | -30 | Y | FO | CW |
| 2017-03-30 | 43 | 24 | | 15 | 30 | Y | FO | CCW |
| 2017-04-04 | 38 | 15 | | 3 | -30 | Y | FO | CW |
| 2017-04-06 | 39 | 13 | | 6 | 30 | Y | FO | CCW |
| 2017-04-07 | 30 | 17 | | 4 | -30 | Y | FO | CW |
| 2017-04-12 | 31 | 13 | | 10 | 30 | Y | FO | CCW |
| 2017-04-13 | 35 | 13 | | 5 | -30 | Y | FO | CW |
| 2017-04-18 | 33 | 14 | | 4 | 30 | Y | FO | CCW |
| 2017-04-19 | 39 | 17 | | 3 | -30 | Y | FO | CW |
| 2017-04-20 | 36 | 17 | | 6 | 30 | Y | FO | CCW |
| 2017-04-26 | 39 | 17 | | 11 | -30 | Y | FO | CW |
| 2017-04-27 | 37 | 22 | 10 | | 30 | Y | FP | CCW |
| 2017-05-10 | 35 | 11 | 11 | | -30 | Y | FP | CW |
| 2017-05-11 | 36 | 10 | 15 | | 30 | Y | FP | CCW |

| | | | | | | | | |
|---|---|---|---|---|---|---|---|---|
| 2017-05-12 | 42 | 26 | 20 | | -30 | Y | FP | CW |
| 2017-05-16 | 41 | 24 | 18 | | 30 | Y | FP | CCW |
| 2017-05-17 | 39 | 13 | 13 | | -30 | Y | FP | CW |
| 2017-05-19 | 35 | 21 | 9 | | 30 | Y | FP | CCW |
| 2017-05-31 | 39 | 28 | 6 | | -30 | Y | FP | CW |
| 2017-06-01 | 39 | 9 | 7 | | 30 | Y | FP | CCW |
| 2017-06-09 | 39 | 14 | 12 | | -30 | Y | FP | CW |
| 2017-06-15 | 49 | 3 | 10 | | 30 | Y | FP | CCW |
| 2019-12-05 | 58 | 36 | | 55 | 30 | Z | FO | CCW |
| 2019-12-06 | 68 | 60 | | 77 | 30 | Z | FO | CCW |
| 2019-12-07 | 55 | 32 | | 59 | 30 | Z | FO | CCW |
| 2019-12-09 | 71 | 37 | | 67 | 30 | Z | FO | CCW |
| 2019-12-12 | 56 | 44 | | 56 | 30 | Z | FO | CCW |
| 2019-12-14 | 57 | 56 | | 60 | 30 | Z | FO | CCW |
| 2020-01-10 | 50 | 56 | | 48 | 30 | Z | FO | CCW |
| 2020-01-21 | 50 | 67 | 26 | | 30 | Z | FP | CCW |
| 2020-01-22 | 49 | 59 | 37 | | 30 | Z | FP | CCW |
| 2020-01-23 | 50 | 65 | 32 | | 30 | Z | FP | CCW |
| 2020-01-24 | 50 | 59 | 39 | | 30 | Z | FP | CCW |
| 2020-01-25 | 65 | 62 | 36 | | 30 | Z | FP | CCW |
| 2020-01-28 | 57 | 50 | | 30 | 30 | Z | FO | CCW |
| 2020-01-29 | 49 | 43 | | 37 | 30 | Z | FO | CCW |
| 2020-01-31 | 53 | 42 | | 39 | 30 | Z | FO | CCW |

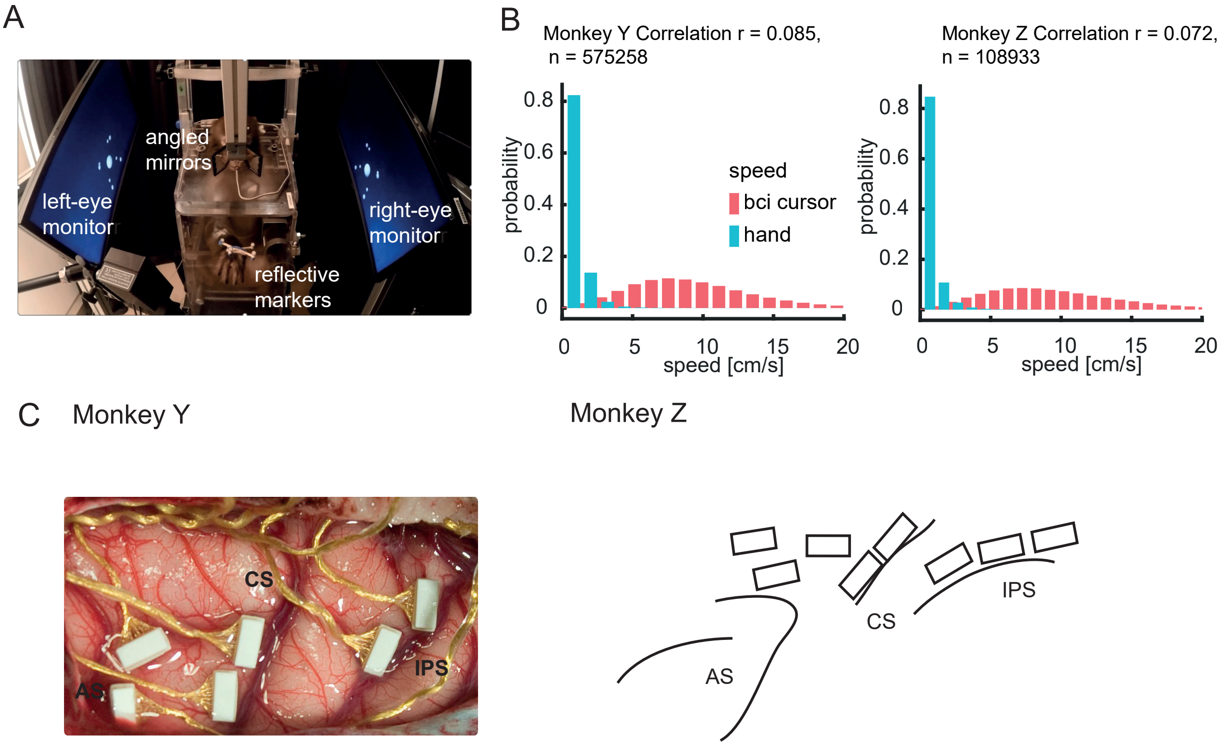
A
angled mirrors
left-eye monitor
right-eye monitor
reflective markers
B
Monkey Y Correlation r = 0.085, n = 575258
Monkey Z Correlation r = 0.072, n = 108933
probability
speed
bci cursor
hand
speed [cm/s]
C
Monkey Y
Monkey Z
CS
IPS
AS

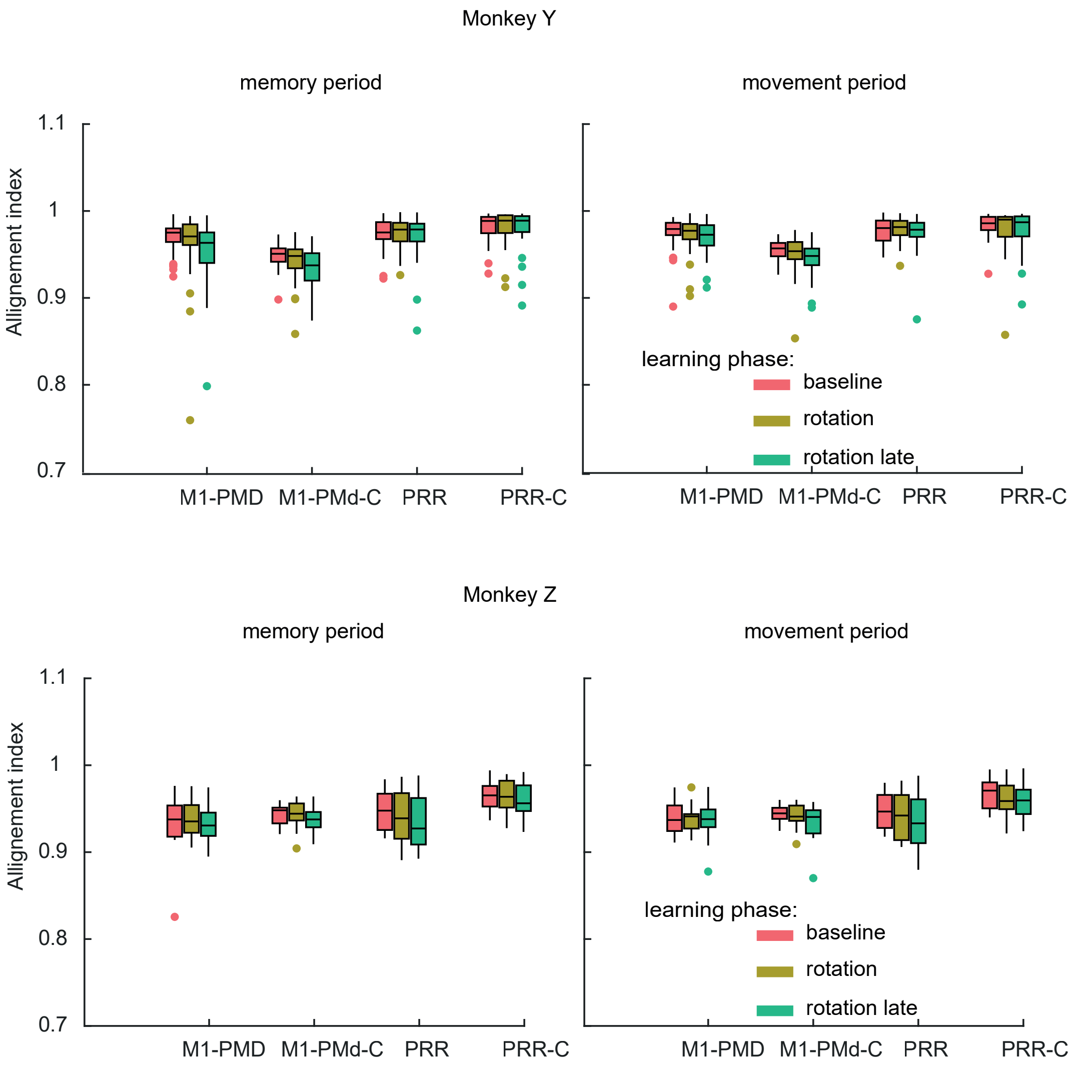
Monkey Y
memory period
movement period
Allignement index
1.1
1
0.9
0.8
0.7
M1-PMD
M1-PMd-C
PRR
PRR-C
learning phase:
baseline
rotation
rotation late
Monkey Z
memory period
movement period
Allignement index
1.1
1
0.9
0.8
0.7
M1-PMD
M1-PMd-C
PRR
PRR-C
learning phase:
baseline
rotation
rotation late

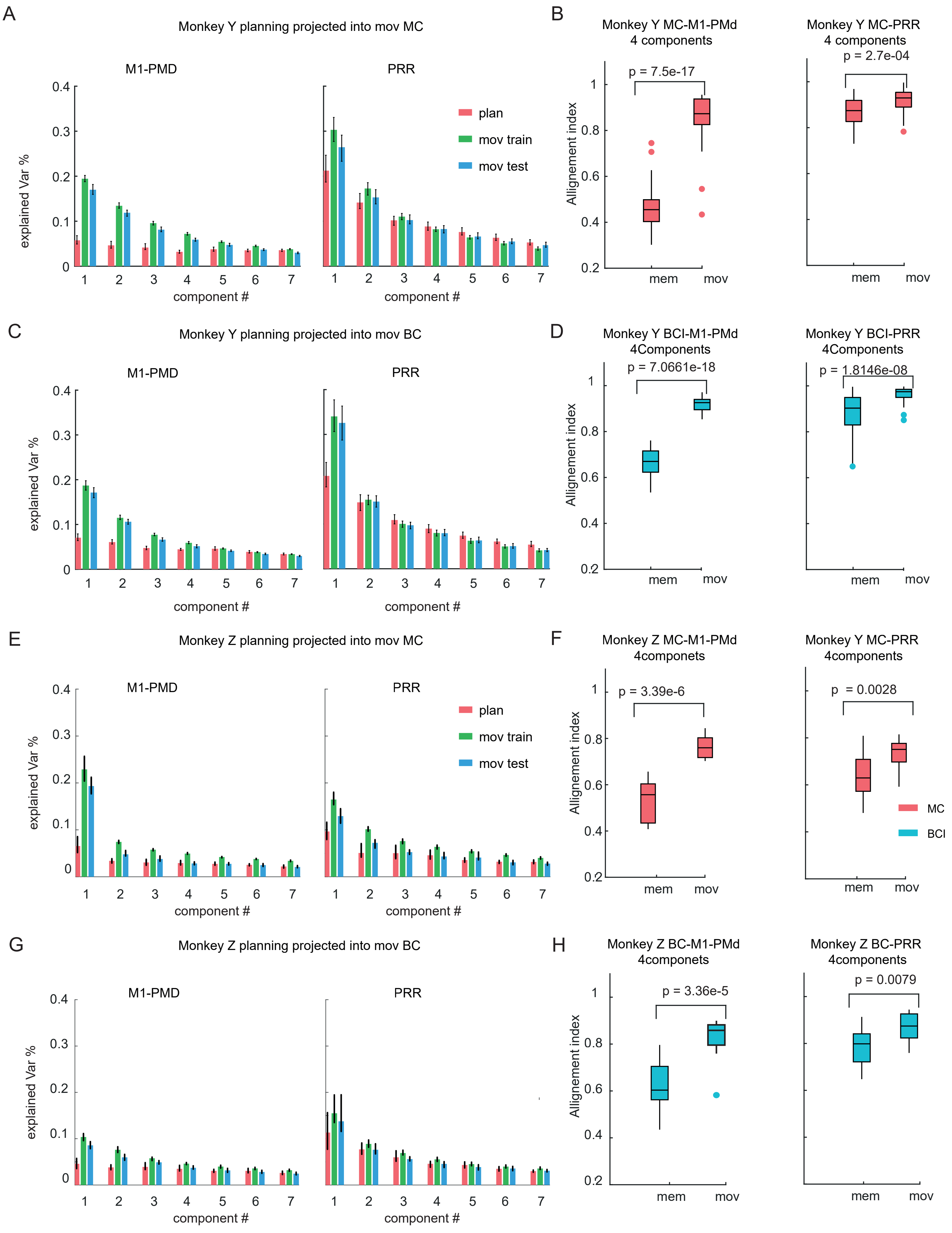
A
Monkey Y planning projected into mov MC
M1-PMD
PRR
plan
mov train
mov test
explained Var %
component #
B
Monkey Y MC-M1-PMd
4 components
p = 7.5e-17
Monkey Y MC-PRR
4 components
p = 2.7e-04
Allignement index
mem
mov
C
Monkey Y planning projected into mov BC
M1-PMD
PRR
explained Var %
component #
D
Monkey Y BCI-M1-PMd
4Components
p = 7.0661e-18
Monkey Y BCI-PRR
4Components
p = 1.8146e-08
Allignement index
mem
mov
E
Monkey Z planning projected into mov MC
M1-PMD
PRR
plan
mov train
mov test
explained Var %
component #
F
Monkey Z MC-M1-PMd
4componets
p = 3.39e-6
Monkey Y MC-PRR
4components
p = 0.0028
Allignement index
MC
BCI
mem
mov
G
Monkey Z planning projected into mov BC
M1-PMD
PRR
explained Var %
component #
H
Monkey Z BC-M1-PMd
4componets
p = 3.36e-5
Monkey Z BC-PRR
4components
p = 0.0079
Allignement index
mem
mov

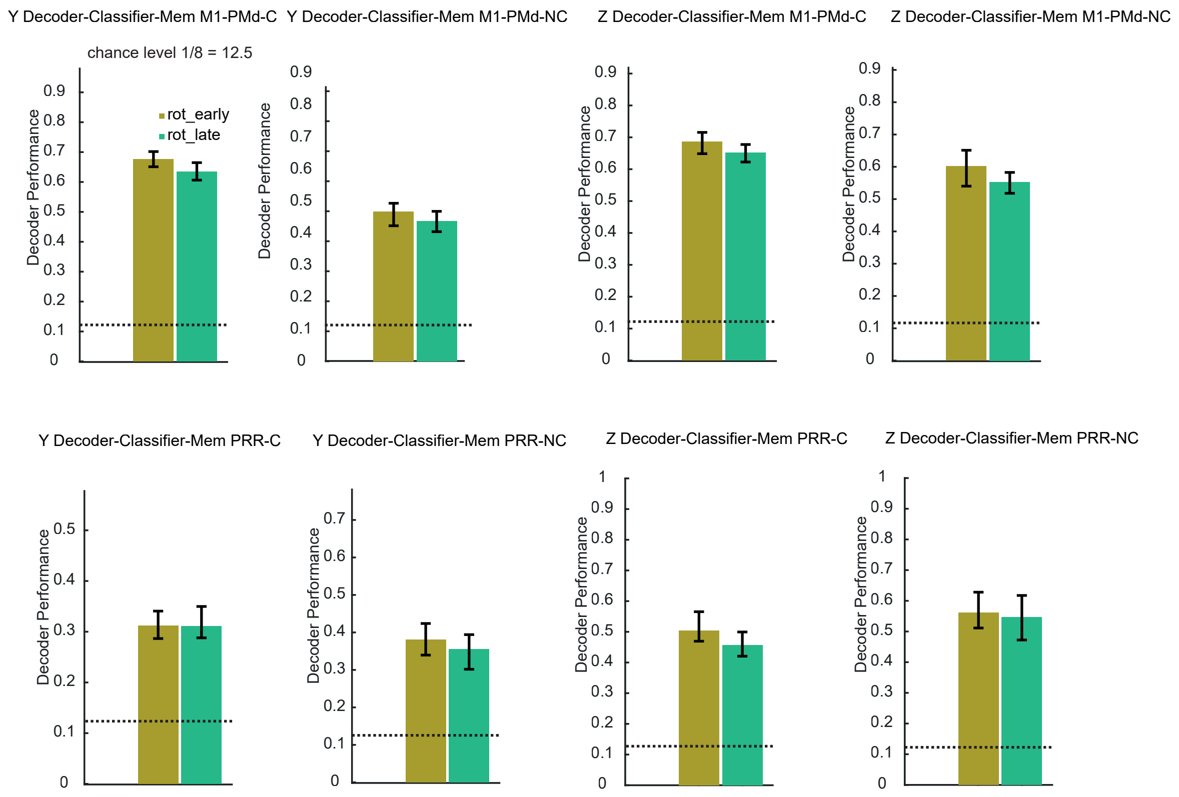
Y Decoder-Classifier-Mem M1-PMd-C
Y Decoder-Classifier-Mem M1-PMd-NC
Z Decoder-Classifier-Mem M1-PMd-C
Z Decoder-Classifier-Mem M1-PMd-NC
chance level 1/8 = 12.5
rot_early
rot_late
Decoder Performance
0
0.1
0.2
0.3
0.4
0.5
0.6
0.7
0.8
0.9
1
Y Decoder-Classifier-Mem PRR-C
Y Decoder-Classifier-Mem PRR-NC
Z Decoder-Classifier-Mem PRR-C
Z Decoder-Classifier-Mem PRR-NC

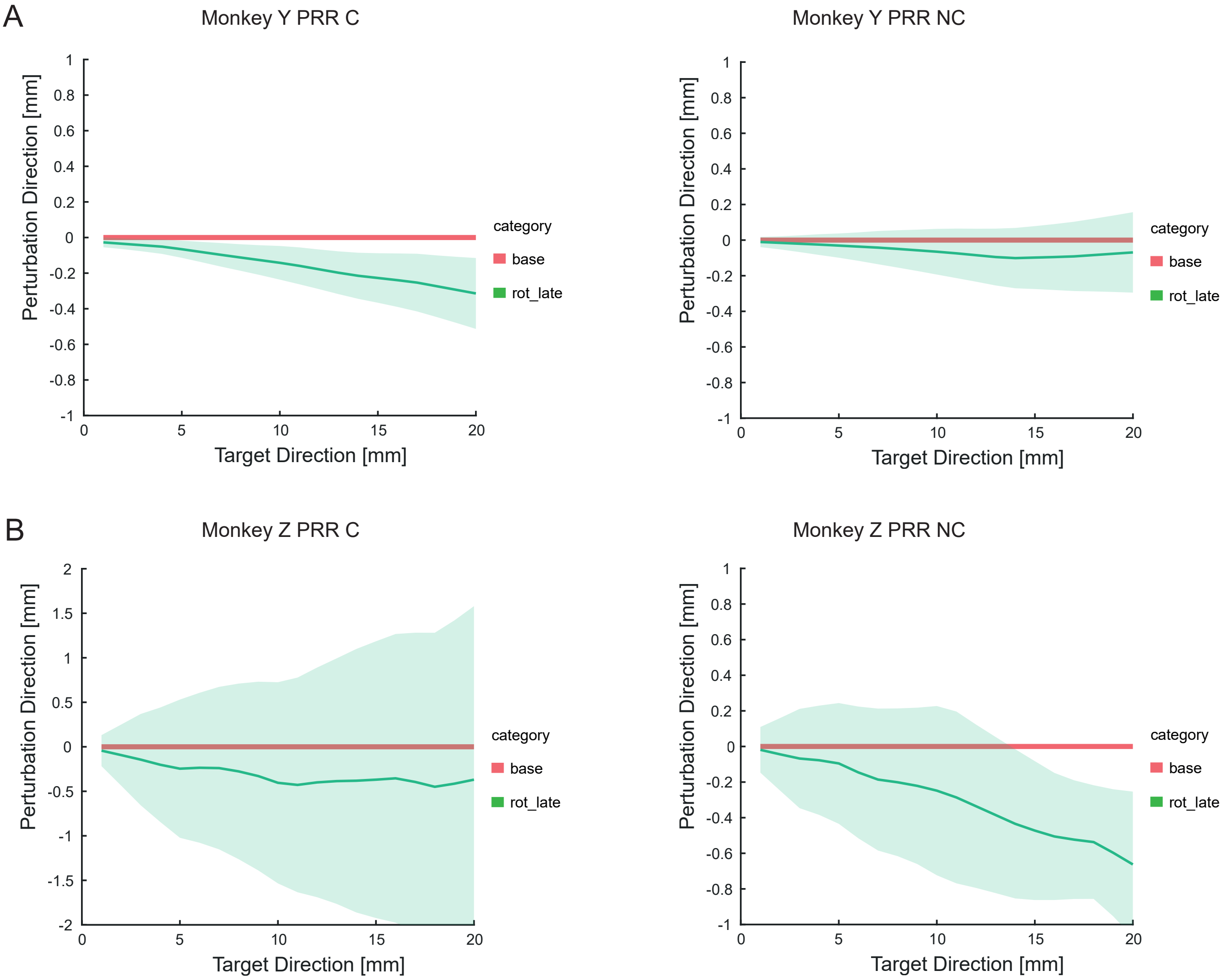
A
Monkey Y PRR C
Monkey Y PRR NC
B
Monkey Z PRR C
Monkey Z PRR NC
Perturbation Direction [mm]
Target Direction [mm]
category
base
rot_late

A

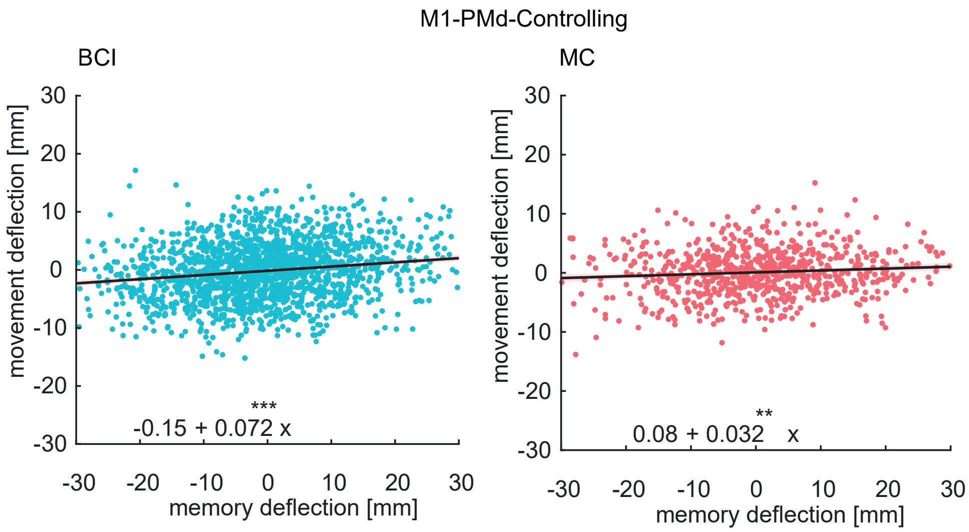

M1-PMd-Controlling
BCI
movement deflection [mm]
30
20
10
0
-10
-20
-30
***
-0.15 + 0.072 x
-30 -20 -10 0 10 20 30
memory deflection [mm]
MC
movement deflection [mm]
30
20
10
0
-10
-20
-30
**
0.08 + 0.032 x
-30 -20 -10 0 10 20 30
memory deflection [mm]
